# Constitutive expression of IκBζ promotes tumor growth and immunotherapy resistance in melanoma

**DOI:** 10.1101/2024.09.19.613946

**Authors:** Antonia Kolb, Ana-Marija Kulis-Mandic, Matthias Klein, Anna Stastny, Maximilian Haist, Beate Weidenthaler-Barth, Tobias Sinnberg, Antje Sucker, Gabriele Allies, Lea Jessica Albrecht, Alpaslan Tasdogan, Andrea Tuettenberg, Henner Stege, Dirk Schadendorf, Stephan Grabbe, Klaus Schulze-Osthoff, Daniela Kramer

**Author notes:** To whom correspondence should be addressed: Prof. Dr. Daniela Kramer, University Medical Center of the Johannes-Gutenberg University of Mainz, Department of Dermatology, Langenbeckstr. 1, 55131 Mainz, Germany.

## Abstract

**Background:** IκBζ, a rather unknown co-regulator of NF-κB, is mostly inducibly expressed and can either activate or repress a specific subset of NF-κB target genes. While its role as a transcriptional regulator of various cytokines and chemokines in immune cells has been revealed, IκBζ’s function in solid cancer remains unclear.

**Methods:** We investigated IκBζ expression in melanoma, and assessed its impact on target gene expression, tumor growth, and response to immunotherapy in melanoma cell lines, mouse models, and patient samples.

**Results:** Unlike in other cell types, IκBζ protein was found to be constitutively expressed in a subfraction of melanoma cell lines, and around 35% of melanoma cases. This atypical expression pattern of IκBζ did not correlate with its mRNA levels or known driver mutations, but instead seemed to result from changes in its post-transcriptional or post-translational regulation. Deleting constitutively expressed IκBζ abrogated the activity and chromatin association of STAT3 and p65, leading to reduced expression of the pro-proliferative cytokines IL-1β and IL-6 in melanoma cells. Consequently, loss of tumor-derived IκBζ suppressed self-sustained melanoma cell growth both *in vitro* and *in vivo*. Additionally, constitutive IκBζ expression suppressed the induction of the chemokines *CXCL9*, *CXCL10*, and *CCL5*, which impaired the recruitment of NK and CD8^+^ T-cells to the tumor, causing resistance to α-PD-1 immunotherapy in mice. Furthermore, the expression of tumor-derived IκBζ also correlated with the absence of CD8^+^ T-cells in human melanoma samples and progressive disease during immunotherapy.

**Conclusion:** We propose that tumor-derived IκBζ could serve as a new therapeutic target and prognostic marker that characterizes melanoma with high tumor cell proliferation, cytotoxic T- and NK-cell exclusion, and unfavorable immunotherapy responses. Targeting IκBζ expression might open up a new therapy option to re-establish the recruitment of cytotoxic cells, thereby resensitizing for immunotherapy.

**Graphical Abstract:** 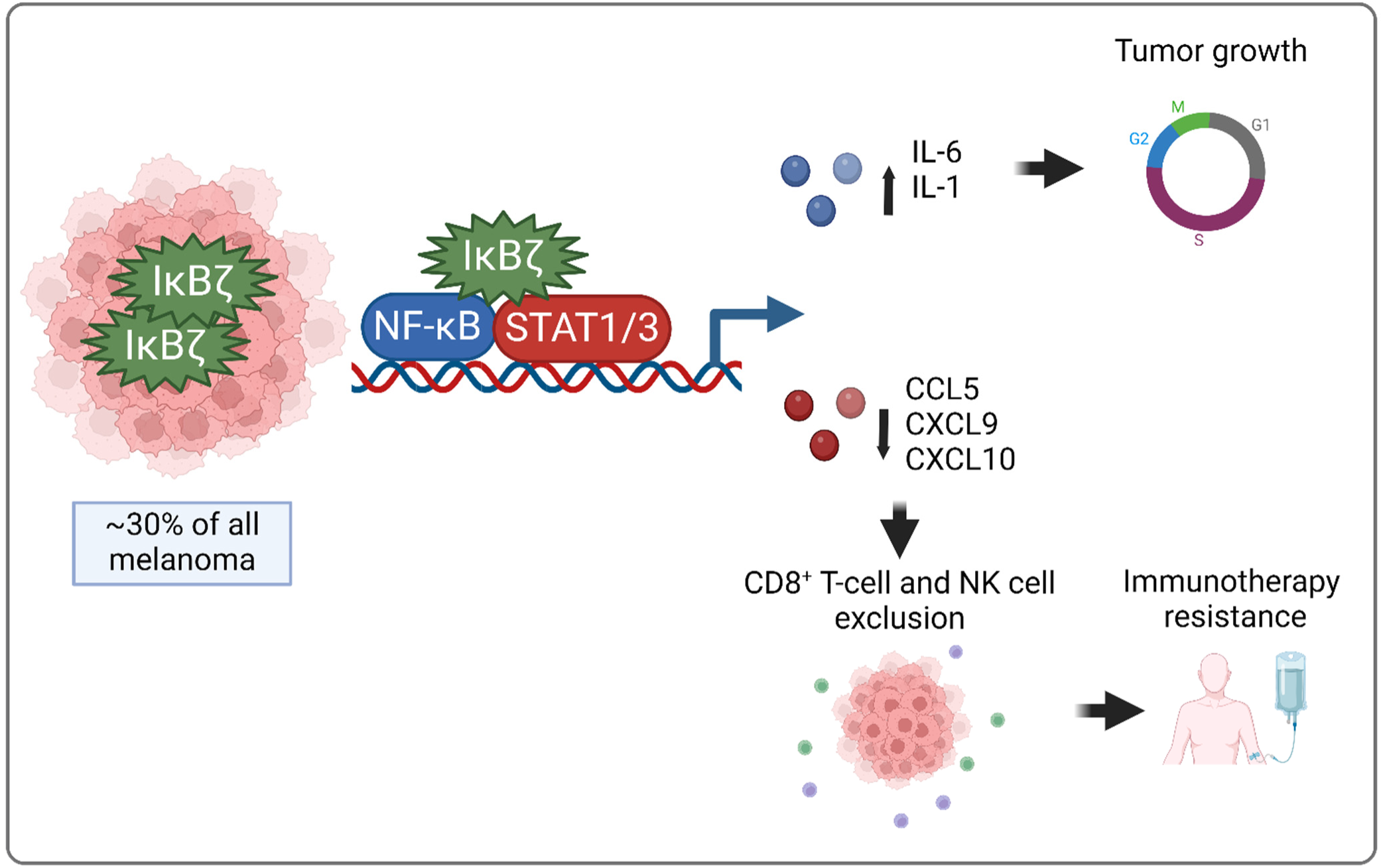

## Introduction

IκBζ, encoded by NFKBIZ, belongs to the NF-κB co-factor family of atypical IκB proteins, which IκBζ is usually not constitutively expressed but rapidly induced upon activation of various immune cells, keratinocytes, or epithelial cells using TLR ligands (e.g. LPS or Imiquimod) or pro-inflammatory cytokines, such as IL-17, IL-1β or IL-36, in an NF-κB- and STAT3-dependent manner ^1^. Depending on the stimulus and cell type, IκBζ protein levels are controlled by post-transcriptional regulation of its mRNA stability through 3’UTR-dependent processing ^2, 3^, or by post-translational modifications, such as phosphorylation or ubiquitination ^4^. Upon protein expression, IκBζ localizes to the nucleus, where it mediates the induction of various pro-inflammatory cytokines and chemokines, such as IL6, IL36, IL8, IL1B, CXCL8, or CXCL2 ^5^. Finally, IκBζ becomes rapidly degraded again to balance the inflammatory response ^2, 3, 5^.

How IκBζ regulates gene expression is still not fully understood. Previously it was shown that IκBζ can interact with transcription factors, such as p50, p52, or STAT3 on the chromatin, thereby possibly inhibiting or increasing their overall activation ^6^. As IκBζ itself lacks any enzymatic activity, it has been suggested that chromatin-bound IκBζ acts as a bridging factor to recruit multiple epigenetic modifiers to chromatin-associated transcription factor complexes, thereby either facilitating or inhibiting the accessibility of specific promoter regions. Up to now, epigenetic modifiers such as TET2, SWI/SNF, or HDAC1 or 2 have been identified as interaction partners of IκBζ, although these interactions seem to be highly variable, dependent on the cell type and stimulus ^7–9^.

While the function of IκBζ in T-cells or keratinocytes, for example, is well established by now ^5, 10^, the role of IκBζ in cancer development and progression has rarely been investigated up to now. So far, only in an aggressive subtype of diffuse large B-cell lymphoma (ABC-DLBCL), the expression and function of IκBζ have been studied in detail ^11^. In this B-cell lymphoma subtype, IκBζ is constitutively expressed, leading to an increased expression of IL-10 and IL-6, which in turn enhance tumor growth and survival.

Besides DLBCL, overexpression of IκBζ has also been detected in other hematological malignancies, including adult T-cell leukemia and mycosis fungoides, the most common subtype of primary, cutaneous T-cell lymphoma. However, the functional consequences of IκBζ expression in these disease contexts remain to be uncovered ^12^. Moreover, the regulation and function of IκBζ expression in solid tumor entities, such as melanoma, are unknown.

To study the role and functional implications of IκBζ expression in solid tumors we chose malignant melanoma, which constitutes a highly immunogenic model tumor, thereby allowing insights into the role of IκBζ for shaping anti-tumor immune responses. Melanoma develops from UV-damaged melanocytes that acquire multiple mutations in proliferation-associated genes, encoding for kinases such as BRAF or NRAS ^13^. Historically, melanoma has been considered a highly lethal disease, as these tumors rapidly proliferate and metastasize to distal lymph nodes and organs. However, the invention of BRAF inhibitors and especially immunotherapy has significantly improved therapy options and overall survival rates for melanoma patients ^14^. Expression of immune-checkpoint proteins such as PD-L1 within the tumor and myeloid-cell compartments constitute a commonly used mechanism of tumor immune evasion that limits effector T-cell responses through activation of pathways involved in T-cell exhaustion, that result in the expression of exhaustion markers including PD-1 and LAG3 ^15^. Application of α-PD-1, α-PD-1L, or α-CTLA4 antibodies block this immune-evasive mechanism, thereby reactivating cytotoxic T-cells in the tumor microenvironment (TME) and subsequently inducing T-cell-dependent tumor cell death. Unfortunately, only 30-45% of melanoma patients respond to initial immunotherapy ^16^. The molecular mechanisms underlying immune checkpoint blockade (ICB) failure remain poorly understood and might be multifactorial. However, as ICB can only reactivate preexisting cytotoxic T-cells in the TME, the exclusion of T-cells and NK cells from the tumor stroma has been identified as one of the major obstacles that contribute to ICB resistance ^17^. Understanding the mechanisms underlying ICB failure might therefore identify new therapeutic targets that could sensitize melanoma patients to immunotherapy treatment approaches.

Here we investigated the expression and function of IκBζ in melanoma, using patient tumor samples, melanoma cell lines, and immunocompetent melanoma mouse models. We observed that a subfraction of melanoma exhibits constitutive IκBζ protein expression, which leads to enhanced STAT3 and NF-κB activation, thereby increasing the expression of various pro-inflammatory cytokines and chemokines. Importantly, IκBζ-dependent gene expression in melanoma sustains and enhances tumor cell proliferation and modulates the cellular organization within the TME. This effect was especially evident for cytotoxic cells, as the constitutive expression of IκBζ in melanoma suppressed their recruitment into the tumor stroma, thus inhibiting α-PD-1 antibody responses *in vivo*. In conclusion, we propose that constitutive IκBζ expression represents an attractive target for new therapy approaches in melanoma, as inhibition of IκBζ can suppress tumor growth and re-store cytotoxic cell infiltration in the TME, thus re-sensitizing tumors for immunotherapy.

## Results

### IκBζ is constitutively expressed in a subgroup of primary and metastatic melanoma

In an initial discovery attempt, we investigated the mRNA and protein levels of IκBζ (encoded by NFKBIZ) in primary and metastatic human and murine melanoma cell lines, harboring different driver mutation profiles. While NFKBIZ mRNA was detectable in varying levels in all investigated melanoma cell lines, a subfraction of cell lines (including human melanoma cell lines LOX-IMVI, SK-MEL-30, and SK-MEL-5, or murine melanoma cell lines D4M-3A and YUMM1.7) displayed constitutive IκBζ protein expression (Figure 1A). Of note, IκBζ protein levels did not correlate with its mRNA expression (Figure 1A), to common mutations in BRAF or NRAS, (Supplementary Figure S1A), or with the activity of IκBζ-associated transcription factors such as NF-κB or STAT3 (Supplementary Figure S1B). Instead, IκBζ protein levels, but not its mRNA levels, were effectively abrogated by treatment with 4EGI-1, an inhibitor of cap-dependent translation (Figure 1B). Vice versa, inhibiting the proteasome with MG-132 restored IκBζ protein expression in melanoma cells that normally do not express IκBζ protein (Figure 1C). Thus, constitutive expression of IκBζ protein in melanoma seems to derive from alterations in its post-transcriptional or post-translational regulation.

**Figure 1.**
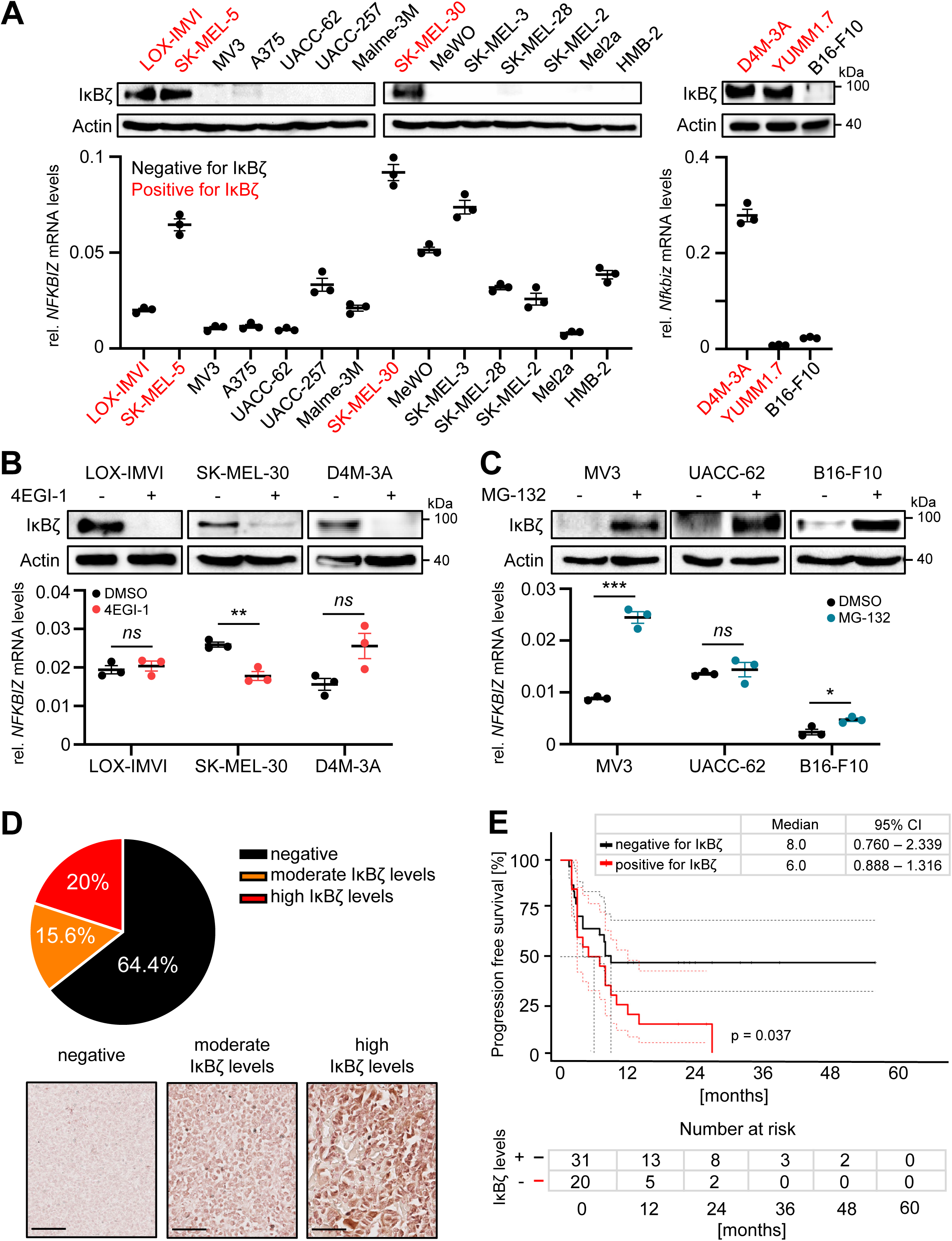
IκBζ protein is constitutively expressed in a subgroup of melanoma cell lines and melanoma patients. **A - C.** β-actin serves as a loading control for immunoblot detection of IκBζ. Human NFKBIZ mRNA expression levels were normalized to RPL37A and murine *Nfkbiz* mRNA expression to *Actb,* respectively. Shown is the mean of 3 biological replicates ± standard deviation (SD). **A.** IκBζ expression levels in various human and murine melanoma cell lines at steady state (Top: immunoblot detection, bottom: mRNA levels). **B.** Detection of IκBζ (NFKBIZ) levels at 24 h after treatment with 25 µM 4EGI-1 or a DMSO vehicle control. **C.** IκBζ expression levels 24 h after treatment with 10 µM MG-132 or a DMSO vehicle control. Shown is the mean of 3 biological replicates ± standard deviation (SD). Significance was calculated using a 2-tailed Student’s t-test (*p < 0.05, **p < 0.01, ***p < 0.001, *ns* = not significant). **D.** Relative amount of IκBζ-positive melanoma samples as detected by immunohistochemistry staining of FFPE samples. *Top:* Analysis of IκBζ protein expression in 90 metastatic melanoma patients using a commercial skin tissue array. *Bottom:* Representative pictures showing no expression of IκBζ, or moderate and high expression of IκBζ in melanoma. Scale bar: 100 µm. **E.** Kaplan-Meier curve showing the progression-free survival (PFS) of 54 melanoma patients grouped according to the presence or absence of tumor-derived IκBζ protein expression. Shown is the median ± 95% CI.

Next, we validated our findings in melanoma patient samples. As mRNA and protein levels of IκBζ do not correlate, assessment of NFKBIZ mRNA levels in patient material was not considered a useful metric to study its constitutive expression. Therefore, we confirmed IκBζ protein expression with immunohistochemistry (IHC) on FFPE samples from primary and metastatic melanoma, using a home-made antibody against human IκBζ, which we intensively validated beforehand (Supplementary Figure S1C and S1D). By analyzing IκBζ protein expression in 90 cases of primary and malignant melanoma, we found that approximately 35% of all melanoma samples displayed IκBζ protein expression, which was uniformly distributed throughout the tumor (Figure 1D). Moreover, we detected gradual changes in expression levels distinguishing moderate (15%) and high (20%) IκBζ expression, although its overall expression intensity did not correlate with tumor stage, or the anatomical site from which the tumor derived (Supplementary Figure S1E and S1F). Interestingly, however, melanoma patients with constitutive IκBζ protein expression in the tumor area showed diminished progression-free survival (Figure 1E).

### Constitutive IκBζ regulates the expression of melanoma-derived cytokines and chemokines

Subsequently, we investigated the functional impact of constitutive IκBζ expression in melanoma. For this purpose, we used shRNA or CRISR-Cas9 to knock down or knock out NFKBIZ (IκBζ) in the IκBζ-expressing LOX-IMVI and D4M-3A melanoma cells (Figure 2A). Global transcriptome analysis of control and NFKBIZ knockdown or *Nfkbiz* knockout cells revealed a significant expression change of 528 genes in LOX-IMVI cells and 267 genes in D4M-3A cells (Figure 2B), which were either up- or downregulated (with a minimum fold change of 2 and a p-value ≤ 0.05). As expected, IκBζ-dependent pathways, that were significantly enriched, comprised the inflammatory response, as well as NF-κB and STAT3 signaling pathways (Figure 2C). Furthermore, we found a significant overlap between IκBζ target genes in melanoma cells and a previously described gene set of inflammatory response-associated hallmarks (Figure 2D and 2E) ^18^. This IκBζ-dependent regulation of inflammatory genes could be validated across various melanoma cell lines, regardless of whether NFKBIZ was knocked down in cells with constitutive expression of IκBζ, or transiently overexpressed in IκBζ negative melanoma cells (Figure 2F and 2G, Supplementary Figure S2). Furthermore, we detected the binding of IκBζ to the respective target gene promoter sites by chromatin immunoprecipitation assays, indicating direct regulation of these target genes by IκBζ (Figure 2H). Thus, we revealed, that constitutively expressed IκBζ in melanoma constitutes a critical and conserved regulator of multiple cytokines and chemokines, including IL6, CXCL8, IL1B, or CXCL10.

**Figure 2.**
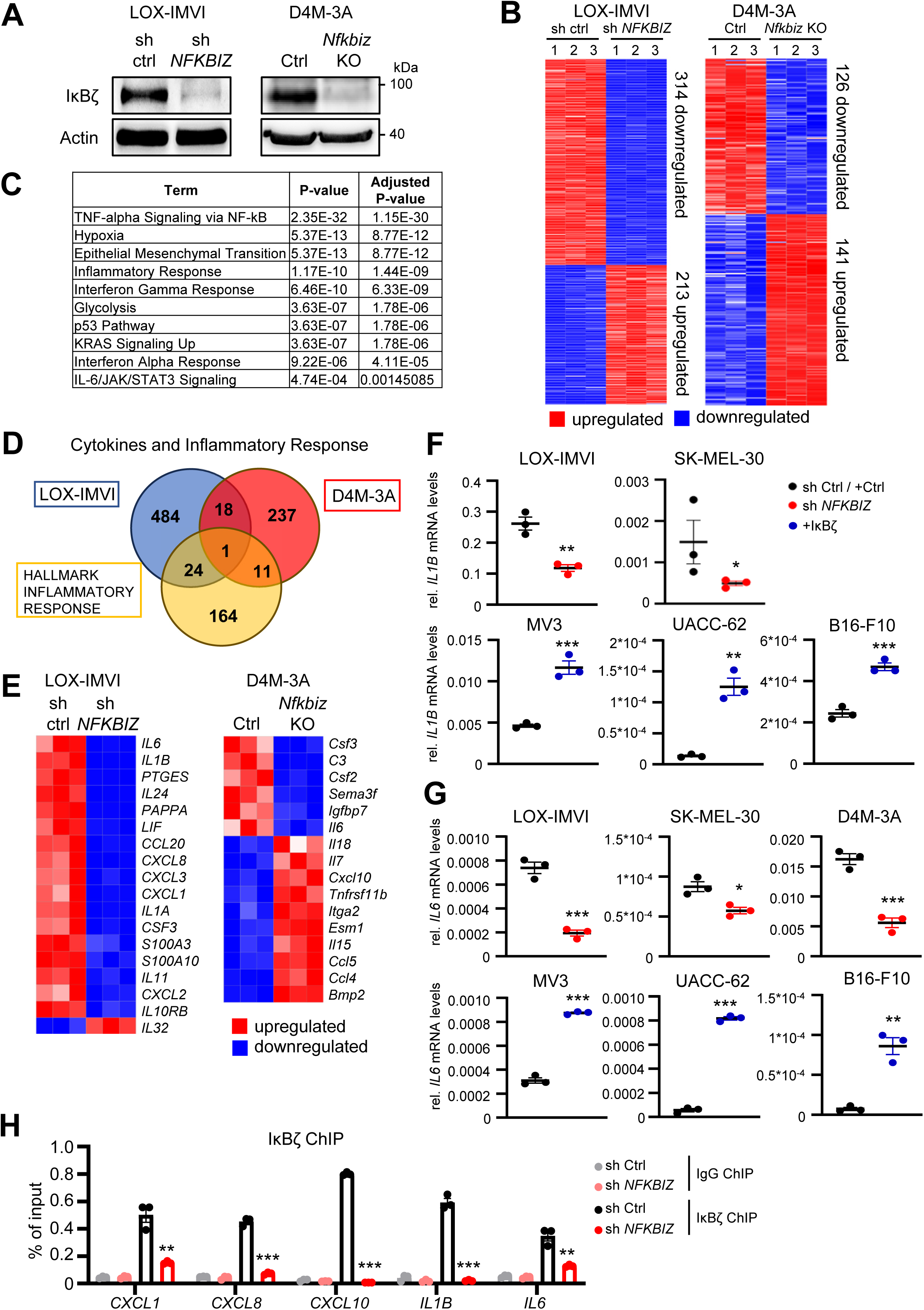
IκBζ regulates the expression of cytokines and chemokines in melanoma cells. **A.** IκBζ knockdown and knockout in LOX-IMVI and D4M-3A cells, respectively. IκBζ was knocked down or deleted by lentiviral transduction using control or NFKBIZ shRNA, or by using an empty control and a *Nfkbiz* CRISPR-Cas9 construct. Efficient knockdown was validated by immunoblot detection of IκBζ, normalized to β-actin. **B.** Heatmaps showing all genes that were either up- (red) or downregulated (blue) upon depletion of IκBζ in LOX-IMVI or D4M-3A cells. Each condition included three biological replicates. IκBζ-regulated genes were defined based on a minimum fold change of 2, and *p* ≤ 0.05 (D4M-3A) or *p* ≤ 0.06 (LOX-IMVI). **C.** Selected results from a gene set enrichment analysis of IκBζ target genes in LOX-IMVI cells using Enrichr (MSigDB Hallmark 2020) ^48^. **D.** Comparison of IκBζ target genes in LOX-IMVI or D4M-3A cells encoding for cytokines and chemokines, with a previously published gene set of inflammatory target genes (GSEA term hallmark inflammatory response) ^18^. **E.** Detailed heatmap of the RNA sequencing results from **B** showing all IκBζ-regulated cytokines and chemokines. **F + G.** Relative expression of IL1B and IL6 in various NFKBIZ knockdown or IκBζ-overexpressing melanoma cells. Relative mRNA levels were normalized to the reference gene RPL37A (human) or *ActB* (murine). **H.** Chromatin immunoprecipitation (ChIP) assays of IgG (as control) and IκBζ in shRNA control and NFKBIZ knockdown LOX-IMVI cells. Data represent the mean of 3 biological replicates ± standard deviation (SD). Significance was calculated using a 2-tailed Student’s t-test (*p < 0.05, **p < 0.01, and ***p < 0.001).

### Melanoma-derived IκBζ promotes self-sustained tumor proliferation *in vitro* and *in vivo*

Next, we investigated the functional consequences of IκBζ and its target gene expression in melanoma. Previous studies suggested, that IκBζ target genes, such as IL6, IL1A, or IL1B, promote tumor cell proliferation, although it is not entirely clear if these cytokines are predominantly made by the tumor itself or the TME ^19–21^. As the expression of these cytokines is strictly dependent on the presence of IκBζ in melanoma, we were interested if IκBζ overexpression or knockdown has a direct effect on the survival or proliferation of the tumor cells. Interestingly, CRISPR-Cas9-mediated knockout of constitutive IκBζ in D4M-3A cells led to decreased cell viability *in vitro* (Figure 3A). Vice versa, overexpression of IκBζ did not change the proliferation and survival of the IκBζ negative cell line MV3 under normal cell culturing conditions. (Supplementary Figure S3A). However, upon withdrawal of FCS, IκBζ overexpression was sufficient to sustain tumor cell proliferation in the non-expressing melanoma cell lines B16-F10 and MV3, while control cells lacking IκBζ stopped proliferation and died (Figure 3B and 3C). Importantly, the proliferation of FCS-depleted control MV3 cells could be restored by the addition of cell culture supernatant from IκBζ-overexpressing MV3 cells, supporting our hypothesis that tumor-derived cytokines drive FCS-independent tumor growth (Figure 3D). Furthermore, supplementation of FCS-depleted cell culture medium with IL-6 and IL-1β, two proliferation-promoting cytokines regulated by IκBζ, was sufficient to re-establish the proliferation of control MV3 cells under starvation conditions (Supplementary Figure S3B).

**Figure 3.**
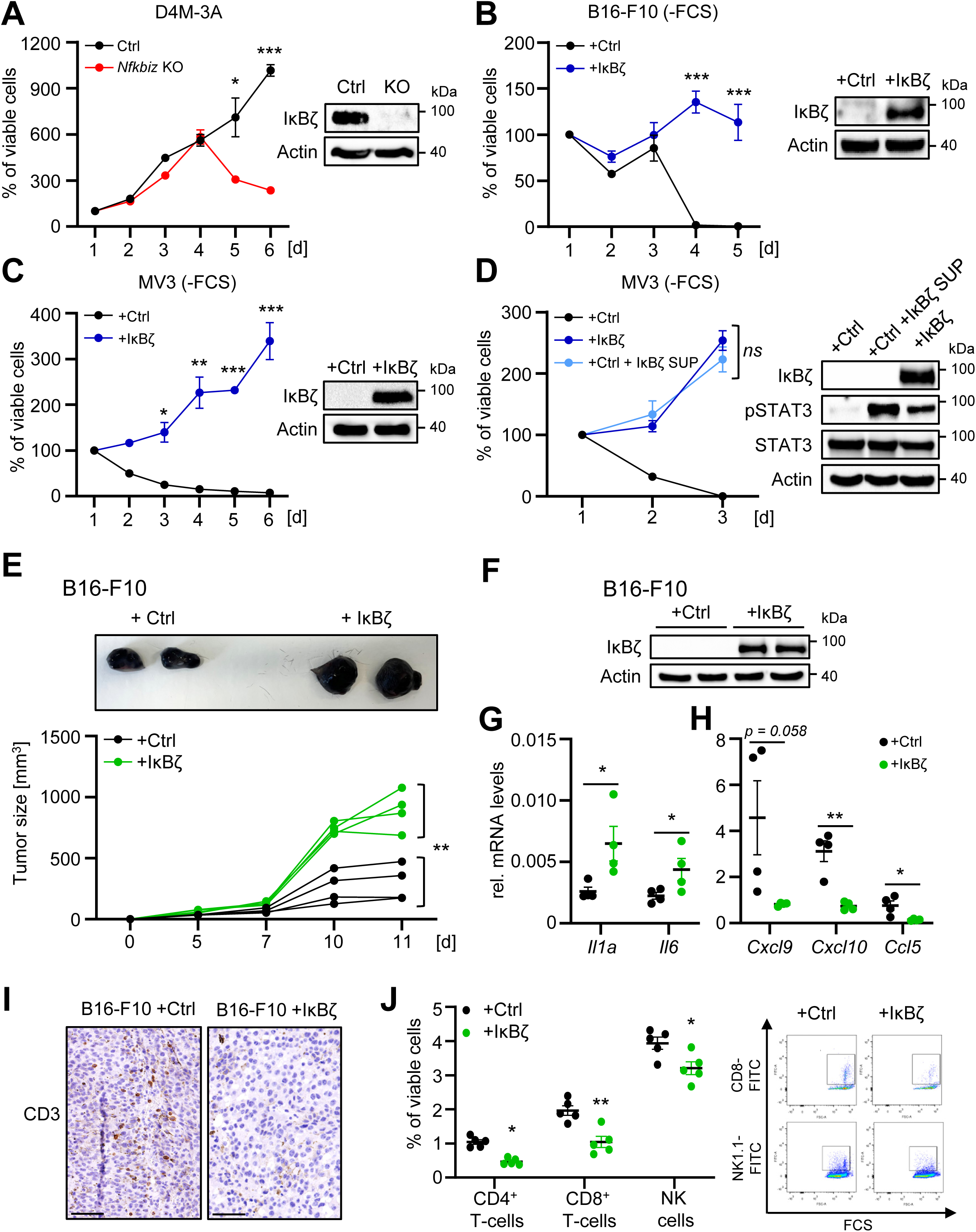
Melanoma-derived IκBζ promotes self-sustained cell proliferation and tumor growth *in vitro* and *in vivo.* **A.- D.** *In vitro* cell proliferation was assessed using the CellTiter-Glo assay (Promega). For calculation of the relative proliferation of cells, values, obtained 24 h after initial seeding, were set to 100%, and follow-up measurements were calculated as the percentage relative to the values from the first time point. Shown are mean values from biological triplicates ± standard deviation. IκBζ overexpression or knockdown were controlled by immunoblotting and normalized to ß-actin for each experiment. **A.** Control or *Nfkbiz* knockout D4M-3A cells cultured in FCS-containing medium. **B.** Control or IκBζ-overexpressing B16-F10 cells, cultured under starvation conditions (without FCS). **C.** Control or IκBζ-overexpressing MV3 cells, cultured under starvation conditions (without FCS). **D.** Same as in **C.** with control cells cultured in the presence of supernatant from IκBζ-overexpressing MV3 cells that were previously cultured without FCS. **E.** Tumor growth of control and IκBζ-overexpressing B16-F10 cells, which were subcutaneously infected in the left and right flank of C57/BL6 mice. Tumor growth was assessed over 11 days until mice with IκBζ-overexpressing tumors had to be sacrificed due to ulcerations. N = 4. **F.** Immunoblot confirming IκBζ overexpression in B16-F10 cells at the endpoint of the experiment. β-actin staining served as a loading control. **G. + H.** Relative gene expression levels of IκBζ target genes in control and IκBζ-overexpressing B16-F10 tumors at the endpoint. N = 4. **G.** Pro-proliferative cytokine expression. **H.** Chemokine expression. Relative mRNA levels were normalized to the reference gene *ActB.* **I.** Immunohistochemical staining of CD3^+^ T-cells in control or IκBζ-overexpressing B16-F10 tumors at the endpoint. Scale: 100 µM. **J.** Flow cytometry analysis of infiltrating T-cells into control or IκBζ-overexpressing B16-F10 tumors. The following markers were applied on living (DAPI negative) cells: cytotoxic CD8^+^ T-cells = CD3^+^ CD8^+^; CD4^+^ T-cells = CD3^+^ CD4^+^ CD25^-^; NK cells = CD3^-^ NK1.1^+^. For *in vitro* experiments, the standard deviation is presented (SD), and for *in vivo* analyses the standard error mean (SEM) is shown. Significance was calculated using a 2-tailed Student’s t-test (*p < 0.05, **p < 0.01, ***p < 0.001, *ns* = not significant).

We then analyzed whether the knockdown or overexpression of IκBζ in melanoma cells also impacted tumor cell proliferation and tumor growth *in vivo*, using the two well-defined murine melanoma cell lines, D4M-3A and B16-F10 ^22, 23^. After subcutaneous injection of control and IκBζ knockout D4M-3A cells, or control and IκBζ-overexpressing B16-F10 cells into immunocompetent C57/BL6 mice, we monitored tumor growth over time. In agreement with our previous *in vitro* results, control D4M-3A cells grew very fast, whereas IκBζ knockout D4M-3A cells initially formed small, palpable tumors, 5-7 days after injection, however, stopped growing at day 7 and later (Supplementary Figure S3C and S3D). Vice versa, overexpression of IκBζ in B16-F10 cells accelerated tumor growth compared to IκBζ negative, control B16-F10 cells, causing premature termination of the experiment on day 11, when IκBζ-overexpressing tumors started to ulcerate (Figure 3E and F). Molecular analysis of the tumors at the endpoint revealed diminished expression of pro-proliferative cytokines in the IκBζ-deleted D4M-3A tumors (Supplementary Figure S3E), whereas IκBζ-overexpressing B16-F10 tumors displayed increased expression of pro-proliferative cytokines, such as *Il6* or *Il1a* (Figure 3G). Thus, IκBζ expression in melanoma promotes tumor cell proliferation by increasing the expression of pro-proliferative cytokines.

Interestingly, we found that IκBζ simultaneously modulated the expression of several chemokines such as *Cxcl9, Cxcl10*, and *Ccl5* in D4M-3A and B16-F10 melanoma *in vivo* (Figure 3H and Supplementary Figure S3E). Therefore, we hypothesized that IκBζ expression in melanoma does not only regulate tumor growth but also shapes the TME. To address this question, we investigated the composition of the TME in tumors of control and IκBζ-overexpressing B16-F10 tumors. Whereas the majority of immune cell subtypes remained unchanged (Supplementary Figure S3F), we detected a reduction of infiltrating CD3^+^ T-cells in IκBζ-overexpressing B16-F10 tumors (Figure 3I). Conversely, knockout of IκBζ increased the numbers of infiltrating CD3^+^ T-cells into D4M-3A tumors (Supplementary Figure S3G). Subsequent detailed analysis by flow cytometry revealed a significant decrease of especially tumor-infiltrating CD8^+^ T-cells and NK cells in IκBζ-overexpressing B16-F10 tumors (Figure 3J). In conclusion, IκBζ expression in melanoma inhibits the infiltration of cytotoxic T-cells into the tumor, probably by suppressing the expression of chemokines such as *Cxcl10, Cxcl9,* and *Ccl5*.

### Constitutive IκBζ expression in melanoma promotes immunotherapy resistance

One important mechanism driving immunotherapy resistance is mediated by the exclusion of cytotoxic T-cells and NK cells from the tumor, due to the downregulation of important chemoattractants, such as CCL5, CXCL9, or CXCL10 ^24–26^. As constitutive IκBζ expression led to both significantly decreased levels of immunoregulatory cytokines and tumor-infiltrating cytotoxic T-cells, we wondered whether IκBζ might contribute to immunotherapy resistance, which is frequently observed in melanoma patients. Analysis of melanoma patients revealed strong IκBζ staining, especially in those patients who failed to respond to immunotherapy with nivolumab, either alone or in combination with ipilimumab (CR = complete response, PD = progression disease) (Figure 4A, Supplementary Figure S4A). This strong IκBζ expression further inversely correlated to the presence of tumor-infiltrating CD8^+^ T-cells (Figure 4B). Based on these findings in human patients, we suggest that constitutive IκBζ expression actively induces immunotherapy resistance by suppressing chemokine expression, thereby inhibiting the recruitment of cytotoxic T-cells into the tumor stroma.

**Figure 4.**
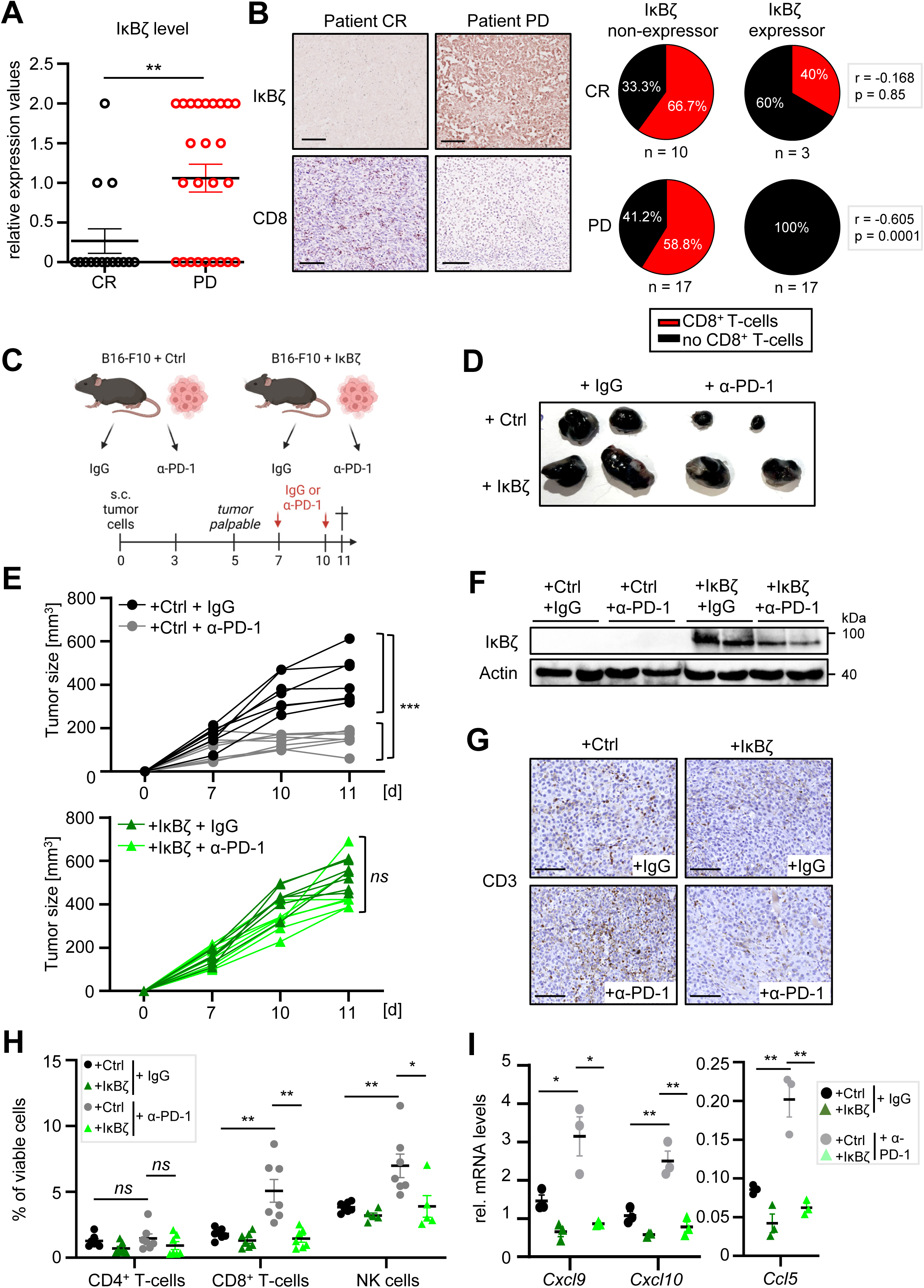
IκBζ expression in melanoma suppresses chemokine expression, leading to diminished T-cell recruitment and increased immunotherapy resistance. **A.** Relative protein levels of tumor cell-derived IκBζ in melanoma samples from patients with different responses to immunotherapy with nivolumab and/or ipilimumab. CR = complete response, PD = progressive disease. No detectable IκBζ expression was graded as 0, moderate expression as 1, and strong expression as 2. B. *Left:* Representative immunohistochemical images demonstrating an inverse correlation of infiltrating CD8^+^ T-cells and IκBζ expression in two melanoma samples, from an immunotherapy-sensitive (CR) and resistant (PD) patient. *Right:* Correlation analysis of tumor-infiltrating CD8^+^ T-cells and IκBζ expression in patients with different immunotherapy responses (CR vs PD). Black = absence of CD8^+^ T-cells, red = presence of CD8^+^ T-cells. **C.** Scheme of the mouse experiment, assessing the response of control and IκBζ-overexpressing B16-F10 tumors to α-PD-1 immunotherapy. Tumor cells were injected in the left and right flank and tumor growth was measured over 11 days. On day 7 and 10, mice received intraperitoneal injections of 200 µg IgG or α-PD-1 antibodies. **D.** Representative picture of the tumors of all 4 experimental groups at day 11. **E.** Tumor growth over time in IgG or α-PD-1 antibody-treated animals with either control B16-F10 tumors (top), or IκBζ-overexpressing B16-F10 tumors (bottom). n = 7 tumors per group. **F.** Immunoblot analysis of IκBζ overexpression in the tumor material at the endpoint, normalized to ß-actin. n = 2 tumors per group. **G.** IHC detection of CD3^+^ T-cells in IgG- or α-PD-1-treated control or IκBζ-overexpressing B16-F10 tumors at the endpoint. Scale: 100 µM. **H.** Flow cytometry analysis of tumor-infiltrating lymphocytes, gated on living cells. The following markers were applied: cytotoxic CD8^+^ T-cells = CD3^+^ CD8^+^; CD4^+^ T-cells = CD3^+^ CD4^+^ CD25^-^; NK cells = CD3^-^ NK1.1^+^. **I.** Gene expression of *Cxcl9*, *Cxcl10,* and *Ccl5* in the tumors at endpoint, normalized to *Actb*. Data represent the mean of 3 biological replicates ± standard error mean (SEM). Significance was calculated using a 2-tailed Student’s t-test (*p < 0.05, **p < 0.01, and ***p < 0.001).

To test this hypothesis, we assessed therapy responses of immunocompetent mice harboring control and IκBζ-overexpressing B16-F10 tumors, which received intraperitoneal injections with IgG or α-PD-1 antibodies (Figure 4C). While control B16-F10 tumors showed constant cell growth inhibition upon treatment with α-PD-1 antibodies, IκBζ-overexpressing B16-F10 tumors retained tumor growth (Figures 4D and 4E). This was not due to changes in the level of IκBζ overexpression, as α-PD-1 antibody treatment did not alter IκBζ expression levels (Figure 4F). Instead, α-PD-1-treatment of control B16-F10 tumors increased the infiltration of CD3^+^ T-cells, especially CD8^+^ T-cells and NK cells, which was not detectable in IκBζ-overexpressing tumors (Figure 4G and 4H, Supplementary Figure S4B and S4C). In agreement, we detected increased expression of *Ccl5*, *Cxcl9,* and *Cxcl10* in α-PD-1 antibody-treated control B16-F10 tumors, whereas IκBζ overexpression in B16-F10 tumors suppressed the expression of both chemokines (Figure 4I). Overall, our data demonstrates that constitutive IκBζ expression in melanoma contributes to increased immunotherapy resistance, supposably by suppressing CCL5, CXCL9, and CXCL10-mediated recruitment of cytotoxic T- and NK-cells into the TME.

### Constitutive IκBζ expression in melanoma enhances the activity and chromatin association of p65 and STAT3 to induce inflammatory gene expression

Finally, we aimed to understand how IκBζ regulates the expression of cytokines and chemokines in melanoma cells. As IκBζ cannot bind DNA directly, it is thought that it interacts with specific transcription factors, such as NF-κB or STAT, on chromatin, thereby changing promoter accessibility by the recruitment of epigenetic modifiers ^1^. NF-κB, STAT1, and STAT3 are key regulators of the previously identified IκBζ target genes, such as IL6, IL1B, CXCL9, and CXCL10. Hence, we closely investigated the activity and function of these transcription factors in IκBζ-expressing and non-expressing melanoma. We first investigated the overall activity of the transcription factors by analyzing the relative phosphorylation levels of STAT1 (Y701), STAT3 (Y705), and IκBα (S32). *In vitro*, knockout of *Nfkbiz* in D4M-3A or knockdown of NFKBIZ in LOX-IMVI cells reduced the phosphorylation levels of STAT3 and STAT1, whereas overexpression of IκBζ in B16-F10 cells increased the STAT3 phosphorylation (Figure 5A). Vice versa, IκBα levels, as a marker for overall NF-κB activity, remained largely unaffected (Supplementary Figure S5A). Moreover, *in vivo* analysis detected reduced phosphorylation levels of STAT3 (Y705) in *Nfkbiz* knockout D4M-3A tumors, whereas IκBζ-overexpressing B16-F10 tumors displayed higher levels of phosphorylated STAT3, thus validating IκBζ-dependent regulation of STAT3 activation *in vivo* (Figure 5B). In agreement with these findings, STAT1, STAT3, and p65 partially localized in LOX-IMVI cells with constitutive IκBζ expression whereas knockdown of NFKBIZ almost completely abolished the association of STAT1, STAT3, and p65 to the chromatin region (Figure 5C). Chromatin immunoprecipitation (ChIP) analyses further revealed specific binding of STAT3 and p65 to the promoter sites of IκBζ target genes (such as IL6 or CXCL10), which was completely lost upon NFKBIZ knockdown (Figure 5D). Together these data indicate that IκBζ-dependent gene expression is potentially mediated by its regulation of STAT3, p65, and STAT1 transcription factor function.

**Figure 5.**
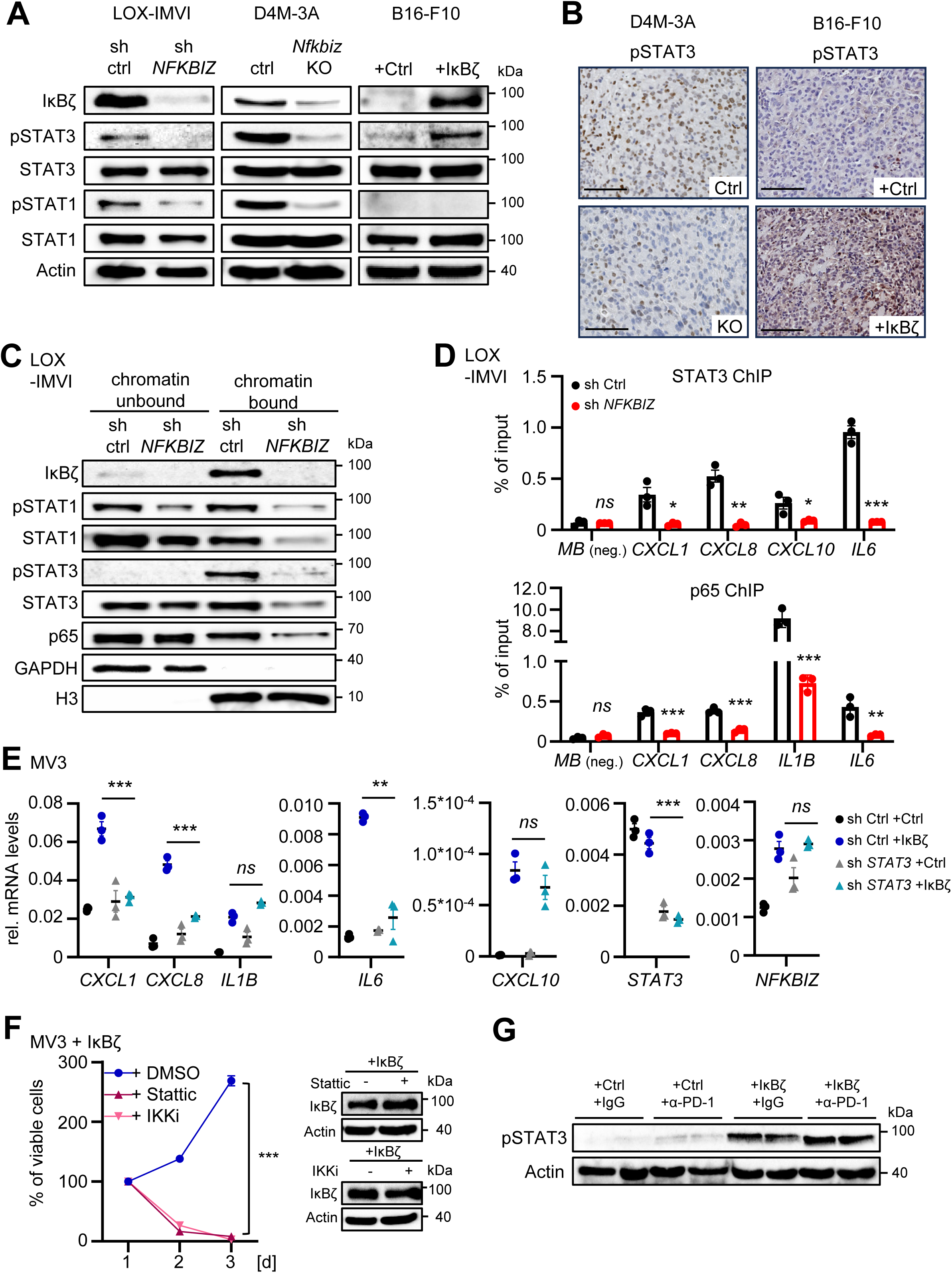
IκBζ regulates the activity and chromatin association of STAT1, STAT3, and p65 in melanoma. **A.** Immunoblot analysis of total and phosphorylated STAT3 (Y705), STAT1 (Y701), IκBα (S32), and p65 (S536) in control or IκBζ-depleted LOX-IMVI or D4M-3A cells, and in control or IκBζ-overexpressing B16-F10 cells. **B.** IHC staining of phosphorylated STAT3 (Y705) in control or IκBζ-depleted D4M-3A tumors, and in control or IκBζ-overexpressing B16-F10 tumors at the experimental endpoint. Scale: 100 µM. **C.** Chromatin fractionation of LOX-IMVI control or NFKBIZ knockdown cells. GAPDH and H3 served as internal controls for the chromatin-unbound and chromatin-bound fraction, respectively. **D.** Chromatin immunoprecipitation of STAT3 and p65 in control or NFKBIZ knockdown LOX-IMVI cells. **E.** Gene expression of IκBζ target genes in control or IκBζ-overexpressing MV3 cells, in the presence or absence of STAT3. STAT3 was lentivirally knocked down using shRNA and control cells were generated using non-coding shRNA (sh ctrl). Gene expression was subsequently analyzed in the presence or absence of transient IκBζ overexpression, normalized to the reference gene RPL37A. **F.** Control or IκBζ-depleted LOX-IMVI cells were treated with 100 ng/mL IL-6 for 24 h. Relative mRNA levels were analyzed and normalized as in (**E**). **G.** Tumor cell viability of MV3 cells, treated with the IKK inhibitor IMD-0354 (5µM) to inhibit NF-κB or a Stattic (10 µM) to suppress STAT1 and STAT3 activity. Cell proliferation was assessed using the CellTiterGlo assay from Promega. N = 3. **H.** Immunoblot detection of pSTAT3 (Y705) levels in IgG- or anti-PD-1-treated control or IκBζ-overexpressing tumors at the endpoint. β-actin controlled equal protein loading. Data represent the mean of 3 biological replicates ± standard deviation (SD). Significance was calculated using a 2-tailed Student’s t-test (*p < 0.05, **p < 0.01, and ***p < 0.001).

To further test this hypothesis, we overexpressed IκBζ in MV3 cells and analyzed IκBζ target gene expression in MV3 cells lacking either STAT3, p65, or STAT1. Consistent with the previous findings, IκBζ overexpression enhanced the expression of several pro-inflammatory genes, such as IL6, CXCL8, and CXCL1, whereas the knockdown of STAT3 abrogated these effects (Figure 5E). Similarly, knockdown of p65 suppressed IκBζ-dependent upregulation of IL1B, IL6, and CXCL8, whereas STAT1 knockdown was sufficient to inhibit IκBζ-dependent upregulation of *CXCL10, IL6, and CXCL8* in MV3 cells (Supplementary Figure S5CBand S5C). Consequently, inhibiting STAT1/3 or NF-κB with Stattic and the IKK inhibitor IMD-0534, respectively, also suppressed self-sustained tumor cell proliferation in MV3 cells (Figure 5F). Furthermore, as STAT3 activation has been implicated in immunotherapy resistance ^27, 28^, we detected elevated phosphorylation and activation of STAT3 in α-PD-1-treated, IκBζ-overexpressing B16-F10 tumors (Figure 5G). In summary, constitutively expressed IκBζ in melanoma stabilizes specific promoter binding and chromatin association of STAT3, p65, and potentially STAT1, depending on the target gene. This leads to increased STAT3 and NF-κB activity, thereby promoting tumor cell proliferation through IL-1 and IL-6, while simultaneously suppressing CCL5, CXCL9, and CXCL10 expression, which impedes T-cell recruitment and contributes to immunotherapy resistance.

## Discussion

IκBζ is a known co-factor of NF-κB, which has previously been found to activate a particular subset of NF-κB target genes, which predominantly encode cytokines, chemokines, and other immune-relevant proteins. While most previous studies focused on the role of IκBζ in immune cells, keratinocytes, or epithelial cells, its role in cancer remains largely unexplored ^5, 10, 29^. Here we investigated the expression and function of IκBζ in melanoma for the first time. Interestingly, we found that a subgroup of melanoma patients and melanoma cell lines displayed constitutive protein expression of IκBζ, which did not correlate with its mRNA expression level. These findings agree with a previous publication, which detected constitutive IκBζ expression in a subtype of diffuse large B-cell lymphoma, called ABC-DLBCL ^11^. Mechanistically, it was shown that constitutive NF-κB activity in ABC-DLBCLs or overactivation of RAS signaling in keratinocytes can result in constitutive IκBζ expression ^11, 30^. However, we could rule out both pathways as drivers of constitutive IκBζ expression in melanoma, as neither the presence of activating BRAF or NRAS mutations nor increased NF-κB activity correlated with constitutive IκBζ protein expression. Instead, inhibition of the eIF4E/eIF4G cap-binding translation complex efficiently suppressed IκBζ protein levels in IκBζ-expressing cells. Vice versa, inhibition of the proteasome restored IκBζ protein levels in non-expressing melanoma cell lines. Therefore, an altered activity or expression of the IκBζ-targeting E3 ubiquitin ligase PDLIM2 ^31^, or known regulators of NFKBIZ mRNA stability, such as Regnase-1 and Regnase-3, could potentially account for the observed constitutive expression levels of IκBζ in melanoma ^2, 3, 32^. Future studies on PDLIM2 or Regnase-1 and -3 might provide insights into the mechanism of constitutive IκBζ expression in melanoma cells.

As IκBζ mainly acts as a transcriptional co-regulator, we investigated target genes of IκBζ by RNA sequencing of IκBζ-deficient LOX-IMVI and D4M-3A cells. Interestingly, IκBζ predominantly regulated the expression of cytokines and chemokines in melanoma. Moreover, we could not only validate the direct binding of IκBζ to the respective target gene promoters but also revealed that the overexpression of IκBζ alone was sufficient to induce target gene expression. Thus, we can rule out indirect signaling events that are needed to trigger IκBζ activation and its target gene expression in melanoma ^33, 34^. Our subsequent analysis revealed that IκBζ promoted the transcription factor functions of STAT3, p65, and STAT1 in melanoma, leading to increased or repressed target gene expression. These results are in agreement with previous studies identifying IκBζ as a direct interaction partner of STAT3 and NF-κB ^35, 36^. Surprisingly, we found that both IκBζ interaction partners could account for IκBζ-dependent target gene activation or repression at the same time. These findings could be explained by the variety of interaction partners that have previously been identified for IκBζ. For example, IκBζ is known to interact with different transcription factor complexes, such as p50 homodimers or p50-p65 heterodimers, leading to gene activation and repression, respectively ^1, 37, 38^. Moreover, previous reports uncovered an interaction of IκBζ with a variety of epigenetic repressors or activators, including HDAC1, HDAC2, TET2, or the SWI/SNF complex ^7–9^. Thus, the presence of different IκBζ interaction partners at specific target gene promoters might explain why IκBζ expression can trigger both, target gene activation and repression.

As a result of IκBζ-dependent target gene expression, we found that IκBζ expression promoted self-sustained tumor cell proliferation and tumor growth. This effect was due to an IκBζ-dependent expression and secretion of IL-1 cytokines and IL-6. These findings align with previous studies, showing that increased or constitutive expression of IL1B, IL1A, and IL6 correlate with accelerated tumor growth and decreased overall survival in melanoma patients ^39–42^. Moreover, IκBζ simultaneously repressed several other chemokines, known to regulate the recruitment of T-cells and NK cells into the tumor microenvironment, such as CXCL9, CXCL10, and CCL5 ^43^. Consequently, we found that melanoma-derived IκBζ expression repressed cytotoxic T-cell or NK-cell recruitment *in vivo*, potentially impairing the effectiveness of immunotherapy by limiting tumor growth inhibition and cell death. Previous reports showed that overall expression of CXCL10 in melanoma mediates the recruitment of cytotoxic T-cells into the tumor stroma ^44^, whereas the re-establishment of CCL5 expression induces NK cell recruitment and melanoma regression ^24^. These results support a model in which IκBζ-dependent repression of *Cxcl10* and *Ccl5* mainly accounts for the observed lack of T-cell infiltration and subsequent immunotherapy resistance, in the α-PD-1 antibody-treated, IκBζ overexpressing B16-F10 melanoma mouse model. Supporting this hypothesis, we also found increased expression levels of *Ccl5*, *Cxcl9,* and *Cxcl10* in the tumors of α-PD-1 antibody-treated control tumors, which was absent in IκBζ-overexpressing B16-F10 tumors treated with α-PD-1 antibodies.

Importantly, a recent study revealed that inhibiting STAT3 with aptamers could re-sensitize melanoma for immunotherapy by upregulating CCL2 and CXCL10 ^27^. Furthermore, NF-κB has previously been reported to upregulate immune checkpoints in cancer, thus contributing to immunotherapy resistance ^28^. In conclusion, we suggest that constitutive IκBζ expression in melanoma increases STAT3 and NF-κB transcription factor activity, subsequently leading to the observed repression in chemokine expression, diminished recruitment of CD8^+^ T-cells and NK cells, and unresponsiveness to immunotherapy. Hence, future studies should investigate whether tumor-derived IκBζ expression correlates with STAT3 and NF-κB activities, not only in melanoma but also in other tumor entities with a frequent constitutive activity of these transcription factors.

## Conclusion

Our study identifies IκBζ as a key factor in melanoma that drives an immunotherapy resistance program coupled with increased tumor growth. We showed that inhibition of IκBζ expression or downstream function represents an attractive new strategy to sensitize melanoma patients to immunotherapy, as its inhibition not only interferes with tumor proliferation but also inhibits two key oncogenic signaling pathways, STAT3 and NF-κB.

## Methods

### Cell culture and treatment

LOX-IMVI, SK-MEL-30, and D4M-3A cells were maintained in RPMI-1640 medium, SK-MEL-3 cells in McCoy’s 5A medium, YUMM1.7 in DMEM/F-12 DMEM (1:1) medium, supplemented with 1% non-essential amino acids, and the other cell lines were maintained in DMEM medium. All media were supplemented with 10% FCS and antibiotics. The cells were grown at 37°C in 5% CO_2_. SK-MEL-2, SK-MEL-5, SK-MEL-28, Malme-3M, UACC-62, and UACC-257 cells were obtained from the NCI-60 human tumor cell line panel. SK-MEL-3 (ACC321) and SK-MEL-30 (ACC151) cells were purchased from the Leibniz Institute DSMZ (Braunschweig, Germany). B16-F10 cells were ordered from ATCC (CRL-6475). LOX-IMVI, MV3, A375, MeWO, Mel2a, and HMB-2 cells were kindly gifted by Matthias Dobbelstein (University of Göttingen, Germany), D4M-3A cells were kindly provided by Tobias Sinnberg (University of Tübingen, Germany). YUMM1.7 cells were kindly provided by Alpaslan Tasdogan (University Hospital Essen, Germany). Recombinant human IL-6 (Cat. 11340064) and IL-1β (Cat. 11340013) were purchased from Immunotools and used at 100 ng/mL final concentration. The following inhibitors were used: 4EGI-1 (Cayman, Cat. 15362), MG-132 (MedChemExpress, Cat. HY-13259), Stattic (Selleckchem, Cat. S7024), and IMD-0354 (Selleckchem, Cat. S2864).

### Lentiviral knockdown, overexpression, or CRISPR-Cas9 mediated knockout of melanoma cell lines

Knockdown cells or overexpressing cells were generated as previously described ^5^. In brief, lentiviral particles were produced in HEK293T cells using the lentiviral vectors pMD2.G (Addgene, Cat.12259) and pCMVR8.74 (Addgene, Cat.22036). For generation of knockdown or overexpressing melanoma cells, the following constructs were used: pLKO.1 (sh ctrl, Addgene, 8453), pLKO.1-TRCN0000147551 (shNFKBIZ, Dharmacon), pLKO.1-TRCN0000020840 (shSTAT3, Addgene), pLKO-TRCN0000280021 (shSTAT1, Sigma), pLKO.1-TRCN0000014686 (shRELA, Sigma) lentiCRISPRv2 pRDI_292 and pRDI_292-NFKBIZ (all kindly provided by Stephan Hailfinger, Münster, Germany) and pAIOsgRNA mCMV-Nfkbiz (GSGM11941-247933929, Dharmacon). All melanoma cells were transduced in the presence of 8 µg/mL polybrene, followed by a selection using 1-2 µg/mL puromycin (InvivoGen).

### Transient overexpression in melanoma cells

Transient overexpression was conducted using the LOX-IMVI Cell Avalanche® Transfection Reagent (EZ Biosystems, EZT-LOX-IMVI). The transfection reagent was mixed with Opti-MEM medium and the appropriate plasmids (pCR3.1 or pCR3-FLAG-NFKBIZ, both kindly provided by Stephan Hailfinger, Münster, Germany). After incubation for 15 min at RT, the DNA complexes were added dropwise to the cells and incubated for 5 h at 37°C and 5% CO_2_. The medium was then replaced by a fresh culture medium, and subsequently, experiments were conducted 48 h post-transfection.

### Mice

All animal experiments were approved by the local animal ethics committee (Landesuntersuchungsamt Rheinland-Pfalz, G22-1-012) and conducted in compliance with German laws and guidelines for animal care. For subcutaneous injection of the tumor cells into the back skin, we used 8-10-weeks old, female C57/BL6 mice (purchased from Charles River). 0.3*10^6^ D4M-3A control or *Nfkbiz* knockout cells, or 0.5*10^6^ B16-F10 control or IκBζ-overexpressing cells were suspended in 100 µL PBS and subcutaneously injected into the left and right back of the animals. When the tumors became palpable, tumor growth was assessed using a caliper, and the tumor volume was calculated using the following formula: tumor volume (mm^3^) = length (mm) x width (mm) x width/2 (mm). For treatment with α-PD-1 antibodies, mice received intraperitoneal injections with 200 µg α-IgG (Leinco Technologies, Cat.I-1177) or α-PD-1 antibodies (Leinco Technologies, Cat.P362), on day 7 and 10, after injection of the tumor cells. Mice were sacrificed on day 11 (B16-F10) or day 18 (D4-M3A) when at least one tumor gained a size of 1000 m^3^ or when the tumors started to ulcerate (as per termination criteria according to the animal study approval).

### Patient material and clinical data

Tissue samples were provided by the University Medical Center Mainz tissue bank and the West German Tumor Centrum, following the regulations of the tissue biobanks and the approval of the local ethics committees (No. 837.226.05(4884) for Mainz and No. 11-4715 for Essen). Clinical characteristics of the patient cohort used to assess IκBζ levels in patients treated with ICB, were stratified by clinical response criteria according to RECIST and are listed in Supplementary Table S3. To detect IκBζ in melanoma samples from different stages and anatomical sites, we used a commercial tissue microarray from BioCat GmbH (Cat. ME1002b-BX). Patients were grouped into 3 subgroups: no detectable IκBζ protein levels (graded as 0), or detectable protein expression levels of IκBζ, ranging between moderate expression (graded as 1), or high overall expression (graded as 2). IκBζ staining deriving in infiltrating immune cells was neglected and excluded from the analysis.

### Flow cytometry

To generate single-cell suspensions, tumors were chopped and digested for 30 min at 37°C with shaking in 1 mL digestion mix (0.25 mg/mL Liberase TM (Sigma, 5401127001), 100 µg/mL DNase I (Sigma, 11284932001), and 0.5 mM CaCl_2_ in RPMI-1640 medium without FCS). After digestion, the cell suspension was passed through a 100 µm cell strainer. Before the antibody incubation, samples were treated with Fc-Block (BioLegend, 101320) for 10 minutes at 4°C and then surfaced-stained for 30 minutes at 4°C with the following antibodies obtained from BioLegend: anti-CD3-APC (Cat. 100236) anti-CD4-PE (Cat. 100408), anti-CD8-FITC (Cat. 100705), anti-Ly6G-PE (Cat. 127608), anti-Ly6C-APC (Cat. 128016), anti-CD11c-PE (Cat. 117308), anti-NK1.1-FITC (Cat. 156507), anti-CD25-PE/Cy7 (Cat. 101915) and anti-CD45-PE (Cat.103106). DAPI (BioLegend, Cat.422801) was used to gate live cells. Data acquisition was performed at the LSRII flow cytometer (Becton Dickinson) with gates set based on the respective isotype controls. Analysis was performed using FlowJo software.

### Histology

After fixation in 4% formaldehyde overnight and paraffin-embedding, 5 μm sections were prepared. Antigen retrieval was performed in 1 mM EDTA (pH 8.0) for 40 min, followed by overnight incubation at 4°C with the following antibodies: anti-phospho-STAT3 (Cell Signaling, Cat. 9145), anti-CD3 (Abcam, Cat. ab16669), anti-CD8 (Agilent, Cat. M7103) or a self-made rabbit antibody against human IκBζ (raised against the peptide CRKGADPSTRNLENEQ, ordered at Eurogentec). After incubation with a peroxidase-coupled secondary antibody, sections were stained with Vector NovaRED (Vector Laboratories, SK-4800) or the SignalStain® DAB Substrate Kit (Cell Signaling, Cat.8059), and counterstained with hematoxylin.

### Cell viability assay

To assess cell viability and proliferation, 1000 – 2000 melanoma cells were seeded on a white 96-well plate per well (PerkinElmer, Cat.6005680) 24 h before the first measurement. In some experiments, control or IκBζ-overexpressing MV3 cells were stimulated with 100 ng/mL of IL-6 or IL-1β 24 h before the first measurement. For treatments with supernatant (SUP), IκBζ-overexpressing MV3 cells were incubated in DMEM medium without FCS for 48 h, after which supernatants were collected. After centrifugation, MV3 control cells were resuspended in the supernatant and seeded for the cell viability assay. During the viability measurement, cells were incubated at 37°C and 5% CO_2_. The CellTiter-Glo® Luminescent Cell Viability Assay (Promega, Cat.G7571) was performed according to the manufacturer’s instructions. The relative luminescence was detected using the Hidex Sense Microplate Reader. Relative luminescence units, (RLU) retrieved from samples 24 h after seeding, were set to 100%, and the relative cell growth was assessed by calculating the fold change in RLUs compared to the 24-h-post-seeding value.

### Immunoblot analysis and co-immunoprecipitation

Immunoblot analysis and Co-immunoprecipitation were performed as described ^45^. The following antibodies were used for immunoblot analysis (all from Cell Signaling): α-IκBζ (Cat. 9244), α-phospho-IκBα (Cat. 2859), α-IκBα (Cat. 9242), α-phospho-STAT1 (Y701, Cat. 9167), α-STAT1 (Cat. 14994), α-phospho-STAT3 (Y705, Cat. 9145), α-STAT3 (Cat. 4904), α-phospho-NF-κB p65 (S536, Cat. 3031) α-NF-κB p65 (Cat. 8242) and α-β-Actin (Cat. 3700). For detection of mouse IκBζ, a self-made antibody was used ^46^. For co-immunoprecipitation of human IκBζ, HEK293T cells were transiently transfected with a pCR3-FLAG-NFKBIZ construct. After cell harvesting, protein lysates were pre-cleared with protein A/G PLUS agarose beads (Santa Cruz, sc-2003) for 1 h at 4°C. The pre-cleared lysates were then incubated overnight at 4°C with antibodies against IκBζ (Cell Signaling, Cat. 9244; Novus Biologicals, Cat. NBP1-89835; or a custom-made antibody from Eurogentec raised against the IκBζ peptide CRKGADPSTRNLENEQ) or against a β-Gal control (Santa Cruz, Cat. sc-19119). The immune complexes were precipitated with protein A/G PLUS agarose beads for 2 h at 4°C, washed with Co-IP buffer, and eluted using 6x SDS-PAGE sample buffer. Pulldown samples were then analyzed by SDS-PAGE and detected by immunoblotting with a FLAG antibody (Sigma, Cat. F1804).

### Chromatin fractionation and chromatin immunoprecipitation

Chromatin fractionation was performed as published before ^47^. α-GAPDH (Cell Signaling, Cat. 2118) and α-histone H3 antibodies (abcam, Cat. ab1791) were used as controls for the chromatin-unbound and chromatin-bound fraction, respectively.

ChIP assays were performed as described ^45^. In brief, chromatin was prepared by crosslinking with 0.25 M DSG (Thermo Fisher, Cat. 20593) for 45 min at RT, followed by cross-linking with 1 % formaldehyde (Thermo Fisher, Cat. 28906). After 20 min of sonification, chromatin was incubated with protein G-coupled Dynabeads (Thermo Fisher, Cat.10003D) and 2 µg self-made antibody against IκBζ (raised against the peptide CRKGADPSTRNLENEQ, ordered at Eurogentec), 3 µg α-STAT3 (Thermo Fisher, Cat. MA1-13042), 2 µg α-p65 (Diagenode, Cat. C15310256), or an IgG antibody as control (Abcam, ab46540), overnight at 4°C on a rotator. ChIP primers for IκBζ target genes are listed in Supplementary Table S2. The primers were self-designed for the promoter regions corresponding to transcription factor-bound sites of STAT3 and p65. Maxima SYBR Green Master Mix (Thermo Fisher, Cat. K0221) was used to perform quantitative PCR. The percentage of input was calculated as described ^45^.

### Gene expression analysis by qPCR and bulk RNA sequencing

The procedure of total RNA extraction and quantitative PCR has been published before ^5^. In brief, total RNA was extracted using Qiazol (Qiagen, Cat. 79306), followed by DNase I digestion to remove contaminating DNA. Afterwards, RNA was either subjected to cDNA synthesis (for qPCR analysis) or library preparation followed by bulk RNA sequencing. Real-time PCR analysis was performed using the CFX384 Touch Real-Time PCR System (Bio-Rad) under the following PCR conditions: initial denaturation 15 minutes at 95°C, followed by 40 cycles of 95°C for 15 seconds and 60°C for 45 seconds. Relative gene expression was quantified by real-time PCR using a Green master mix (Genaxxon, Cat. M3023) and self-designed primers (Supplementary Table S1). For calculation of the relative mRNA levels, the target genes’ Ct values were normalized to the Ct values of the reference genes RPL37A for human samples and *ActB* or *Hrpt1* for murine samples using the 2-ΔCT method.

For transcriptome analysis, both sequencing and library construction from total RNA were performed by Novogene (Munich, Germany) in the case of D4M-3A cells, or by the Genomic Core Facility (Münster, Germany) in the case of LOX IMVI cells. For RNAseq analysis of the human samples, libraries were constructed with the Ultra RNA Library Prep Kit and sequencing was performed using the Illumina NextSeq High Output kit. For analysis of the D4M-3A cells, quantified libraries were sequenced on Illumina Novaseq 6000 platform with paired-end 150 cycle (PE150) sequencing, according to effective library concentration and data amount. Sequencing strategy was paired-end 150 cycles (PE150). CLC Genomics workbench (v24.0; Qiagen) was used to further process the sequencing raw reads. Mapping against the human reference genome hg38 (GRCh38.109) or the mouse reference genome mg39 (GRCm39) was performed by the NGS core facility (Research Center for Immunotherapy) using CLC’s default settings (mismatch cost = 2; insertion cost = 3; deletion cost = 3; length fraction = 0,8; similarity fraction = 0,8; global alignment = No; strand-specific = both; library type = bulk; maximum number of hits for a read = 10; count paired reads as two = no; ignore broken pairs = yes). Differentially expressed genes were filtered for a minimum absolute fold change > 2 with a difference cut-off of absolute >4 and an adjusted p-value ≤ 0.05, between control and NFKBIZ knockdown or knockout cells. Raw data of the gene expression data sets were published under the GEO accession number xxx. Heatmaps were generated using Morpheus software (Morpheus, https://software.broadinstitute.org/morpheus).

## Statistics

For all experiments, at least 3 independent biological replicates were generated and analyzed. Subsequently, significance was calculated using a 2-tailed Student’s t-test (*p < 0.05, **p < 0.01, and ***p < 0.001).

## Abbreviations

TME: tumor microenvironment
ICB: immune checkpoint blockade
IHC: Immunohistochemistry
ChIP: Chromatin immunoprecipitation
CR: complete response
PD: progressive disease
IKKi: IKK inhibitor
STAT: Signal Transducers and Activators of Transcription
SUP: Supernatant

## Acknowledgments

The study was supported by grants from the Foundation of Experimental Biomedicine (D.K.), and the Deutsche Forschungsgemeinschaft TRR156/3 (project number 246807620) (D.K. and S.G.), and TRR355/1 (project number 490846870) (D.K.). We thank Claudia Braun from the Histology Core Facility (University Medical Center Johannes Gutenberg-University Mainz), Bonny Adami, and Dagmar Löck (University Medical Center Johannes Gutenberg-University Mainz) for experimental support. Human biological samples and related data were provided by the Westdeutsche Biobank Essen (WBE/SCABIO, University Hospital Essen, University of Duisburg-Essen, Essen, Germany; approval no. 11-4715) and the Tissue Biobank of the University Medical Center Mainz, approval no. 837.226.05(4884). The graphical abstract and scheme in Figure 4 were created with Biorender (Kramer, D. (2023) BioRender.com/k18x269 and BioRender.com/e11y470).

## Author contributions

A.K., A.M.K.-M., A.S., performed experiments and data analysis. M.K. analyzed the RNA sequencing data. T.S. provided D4M-3A cells, K.S.-O. donated cells from the NCI 60 panel, and T.S. helped planning the mouse experiments. B.W., H.S., M.H., A.T., G.A., A.S., L.J.A., D.S., and S.G. selected the patient samples and helped in interpreting the human data. A.K. and D.K. drafted the first version of the manuscript. All authors reviewed provided feedback and approved the manuscript. All authors read and approved the manuscript.

## Ethical statement

All animal experiments were approved by the local animal ethics committee (Landesuntersuchungsamt Rheinland-Pfalz; G22-01-012). All experiments involving patient material were approved by the local ethics committee of Mainz (No. 837.226.05(4884)) and Essen (No. 11-4715).

## Additional information

**Supplementary Figure S1.** Correlation of IκBζ expression with key driver mutations and transcription factor activation, and additional information of patient samples.

**Supplementary Figure S2.** IκBζ target gene expression in additional melanoma cell lines.

**Supplementary Figure S3.** Additional data on IκBζ-dependent tumor cell proliferation and tumor growth.

**Supplementary Figure S4.** Gating strategy and FACS plots from the analysis of α-PD-1 antibodies-treated control and IκBζ-overexpressing B16-F10 tumors.

**Supplementary Figure S5.** Additional data on IκBζ-dependent regulation of the transcription factor function of STAT1, STAT3, and p65.

**Supplementary Table S1.** Gene expression primer sequences.

**Supplementary Table S2.** ChIP primer sequences.

**Supplementary Table S3.** Summary of the clinical characteristics of the melanoma patients.

