## Supplementary Information for "Constitutive expression of IκBζ promotes tumor growth and immunotherapy resistance in melanoma"

### Additional information

**Supplementary Figure S1.** Correlation of IκBζ expression with key driver mutations, transcription factor activation, and additional information of patient samples.

**Supplementary Figure S2.** IκBζ target gene expression in additional melanoma cell lines.

**Supplementary Figure S3.** Additional data on IκBζ-dependent tumor cell proliferation and tumor growth.

**Supplementary Figure S4.** Gating strategy and FACS plots from the analysis of α-PD-1 antibody-treated control and IκBζ-overexpressing B16-F10 tumors.

**Supplementary Figure S5.** Additional data on IκBζ-dependent regulation of the transcription factor function of STAT1, STAT3, and p65.

**Supplementary Table S1.** Gene expression primer sequences.

**Supplementary Table S2.** ChIP primer sequences.

**Supplementary Table S3.** Summary of the clinical characteristics of human melanoma patients.

**A**

| | Cell line | Primary vs metastasis | *BRAF* | *TP53* | *TERT* | *CDKN2A* | Other mutations |
|---|---|---|---|---|---|---|---|
| IκBζ-expressing cells | **LOX-IMVI** | metastasis | yes | no | yes | yes | *CCND1, NF1* |
| | **SK-MEL-5** | metastasis | yes | no | yes | yes | |
| | **SK-MEL-30** | primary | yes | yes | yes | yes | *NRAS* |
| | **D4M-3A** | primary | yes | no | no | no | *Pten* |
| | **YUMM1.7** | primary | yes | no | no | yes | *Pten* |
| IκBζ non-expressing cells | MV3 | metastasis | no | no | no | no | |
| | MeWO | metastasis | no | yes | yes | yes | *NF1, MAPK3* |
| | A375 | primary | yes | no | yes | yes | |
| | UACC-62 | primary | yes | no | no | yes | *PTEN* |
| | UACC-257 | primary | yes | no | yes | yes | *CDK4* |
| | Malme-3M | metastasis | yes | no | yes | yes | |
| | SK-MEL-3 | metastasis | yes | yes | no | yes | |
| | SK-MEL-2 | metastasis | no | yes | yes | no | *NRAS* |
| | SK-MEL-28 | primary | yes | yes | yes | no | *NRAS, PTEN, CDK4* |
| | Mel2a | metastasis | yes | no | no | no | |
| | HMB-2 | metastasis | yes | no | no | no | |
| | B16-F10 | primary | no | no | no | yes | |

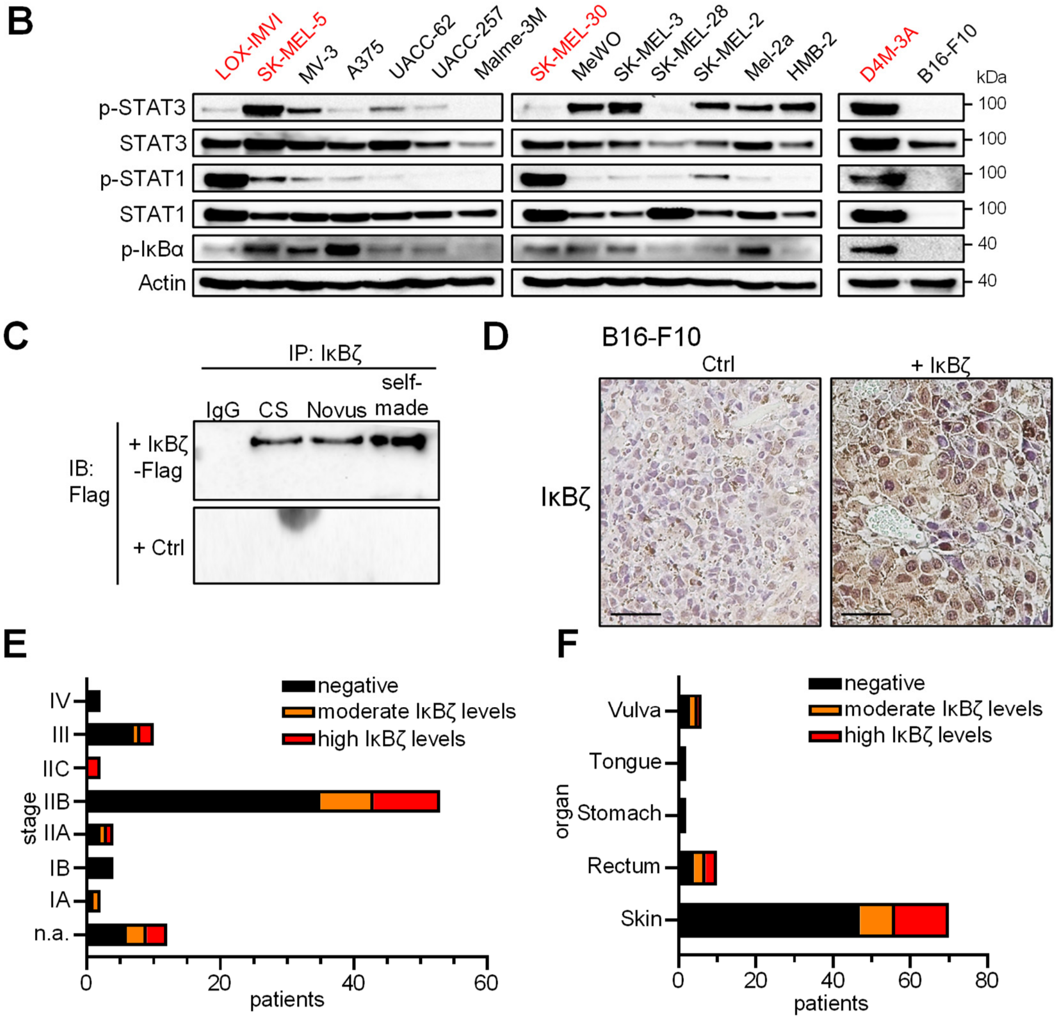


**Supplementary Figure S1. Correlation of IκBζ expression with key driver mutations, transcription factor activation, and additional information of patient samples. A.** Genomic characteristics of all used melanoma cell lines. **B.** Immunoblot analysis of pSTAT3 (Y705), pSTAT1 (Y701), and pIκBα (S32) in all investigated melanoma cell lines at steady-state. **C.** Co-immunoprecipitation of human IκBζ in transfected HEK 293T cells. Cells were transiently transfected with a control plasmid or flag-tagged IκBζ. For immunoprecipitation (IP) of IκBζ, several antibodies were applied (CS = α-IκBζ from Cell Signaling; Novus = α-IκBζ from Novus; self-made = self-made antibody raised against human IκBζ). Immunoblot detection (IB) was performed using a flag antibody. **D.** IHC staining of human IκBζ in B16-F10 tumors harboring expression plasmids for an empty control (Ctrl) or a human IκBζ. Scale: 50 µm. **E. + F.** Characteristics of the patient samples used for Figure 1. **E.** Stage of the disease. **F.** Anatomical tumor site.

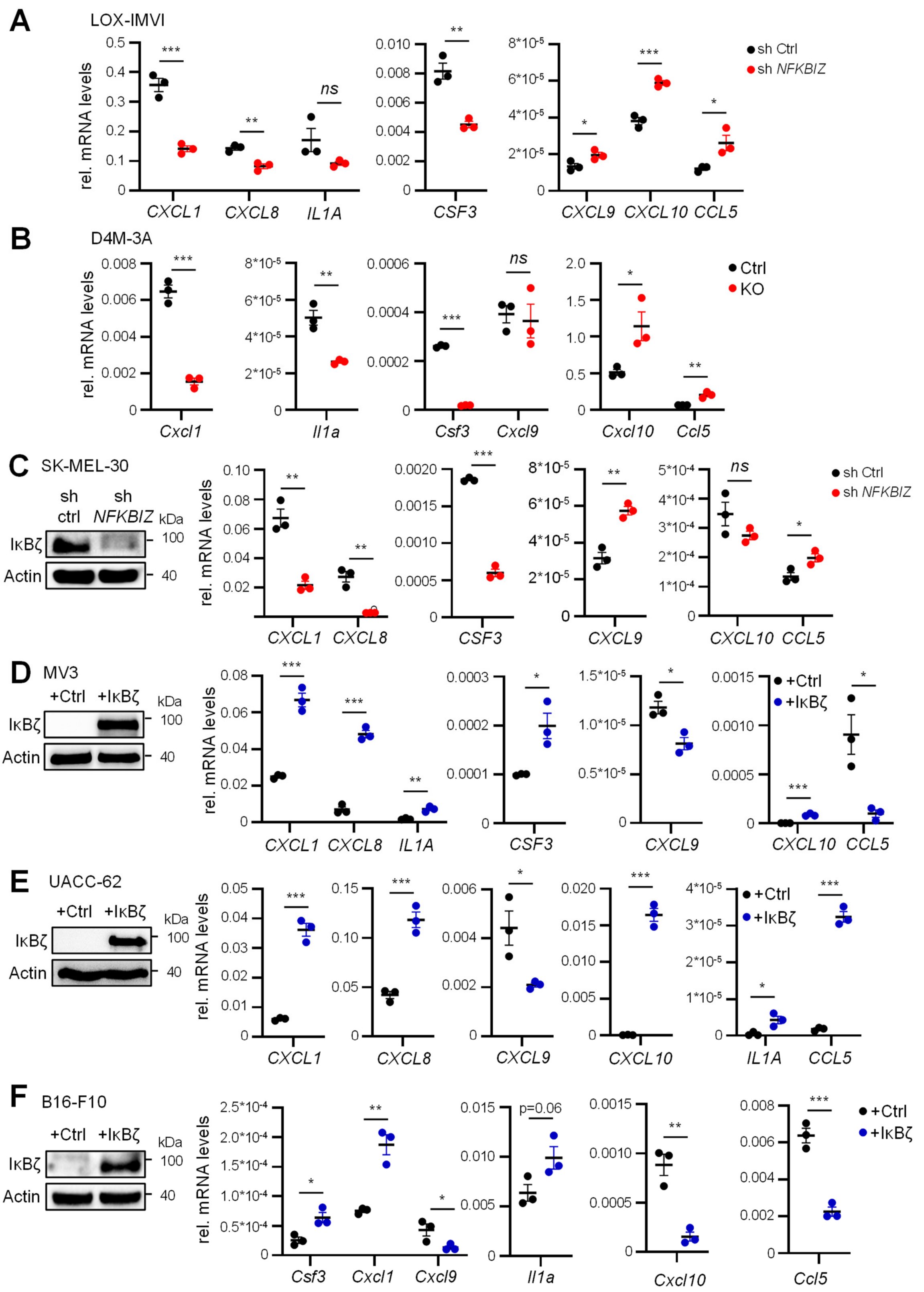


**Supplementary Figure S2. IκBζ target gene expression in additional melanoma cell lines**. *RPL37A* was used for normalization of all human gene expression data, and *Actb* was used to normalize gene expression data of murine cells. β-actin served as a loading control for immunoblot analysis. **A.** Validation of IκBζ target genes in control and *NFKBIZ* knockdown LOX-IMVI cells. **B.** IκBζ target genes in ctrl and KO D4M-3A cells. **C.-F.** Knockdown or overexpression of IκBζ in various melanoma cell lines, and subsequent analysis of IκBζ target gene expression. **C.** SK-MEL-30 cells. **D.** MV3 cells. **E.** UACC-62 cells. **F.** B16-F10 cells. Shown is the mean of 3 biological replicates ± standard deviation (SD). Significance was calculated using a 2-tailed Student's t-test (*p < 0.05, **p < 0.01, ***p < 0.001, *ns* = not significant).

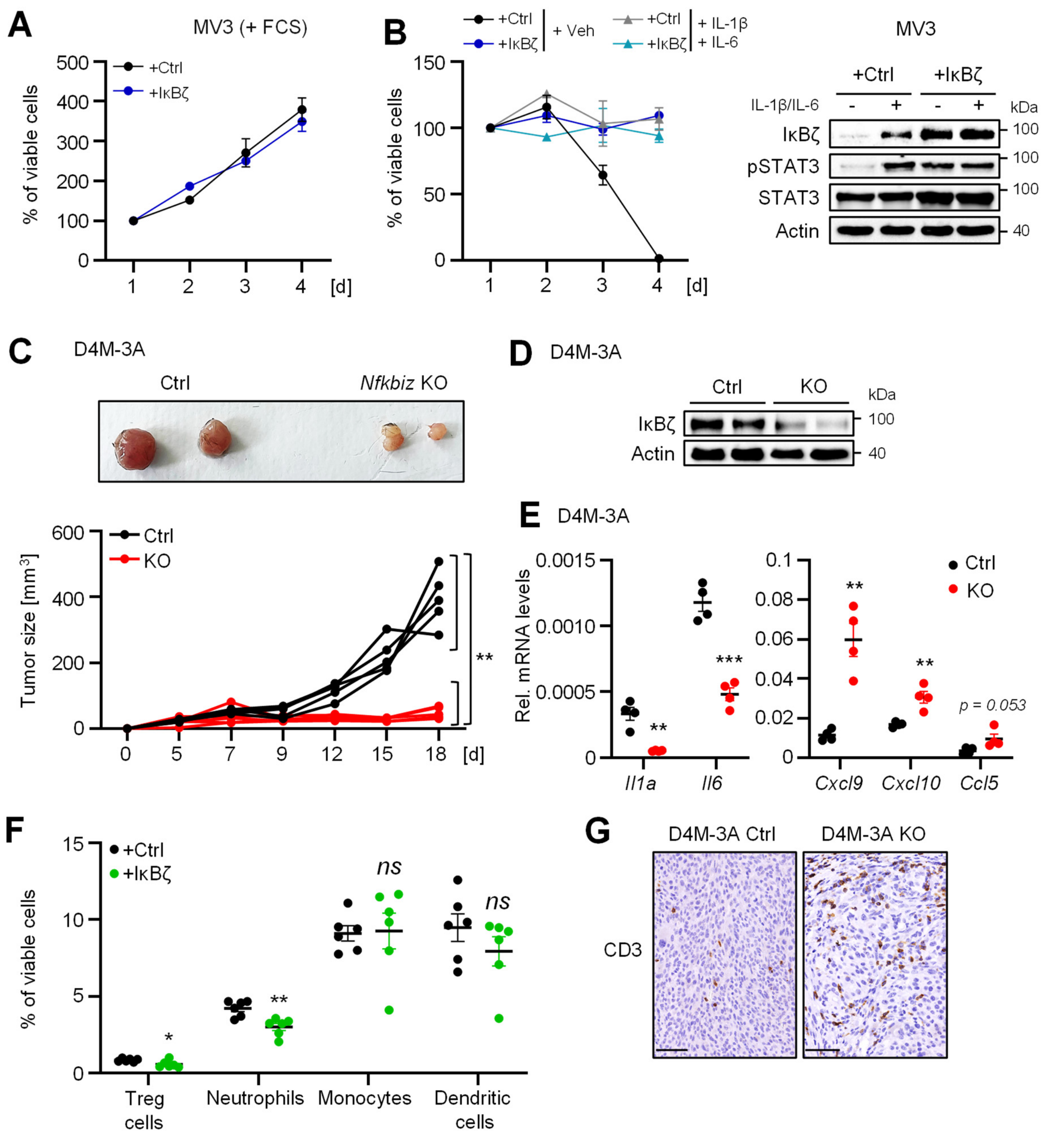


**Supplementary Figure S3. Additional data on IκBζ-dependent tumor cell proliferation and tumor growth. A. and B.** The calculation of the relative proliferation of the cells was done as described in **Figure 3. A.** Control or IκBζ-overexpressing MV3 cells, cultured in complete DMEM medium supplemented with 10 % FCS. **B.** Same cells as in **A.** but cultured under starvation conditions (without FCS). Additionally, cells were treated with 100 ng/mL IL-1β and 100 ng/mL IL-6. IκBζ overexpression was controlled by immunoblotting, normalized to β-actin levels. **C.** Tumor growth of control and IκBζ-depleted D4M-3A cells, which were subcutaneously injected in the left and right flank of C57/BL6 mice. Tumor growth was assessed over 18 days. N = 5. **D.** IκBζ protein levels of tumors at the endpoint were analyzed by immunoblotting. β-actin staining serves as a loading control. **E.** Relative gene expression levels of IκBζ target genes in control and IκBζ-depleted D4M-3A tumors at the endpoint. N = 4. Relative mRNA levels were normalized to the reference gene *ActB.* **F.** Flow cytometry analysis of infiltrating immune cells into control or IκBζ-overexpressing B16-F10 tumors. The following markers were applied on living (DAPI negative) cells: regulatory T-cells = CD3+, CD4+, CD25+, Neutrophils = Ly6G+, Monocytes = Ly6C+, and dendritic cells = CD11c+. N = 5-6. **G.** Immunohistochemical staining of $CD3^+$ T-cells in control or IκBζ-depleted D4M-3A tumors at the endpoint. Scale: 100 µm. Data represent the mean ± standard deviation (SD) for *in vitro* experiments, and the standard error (SEM) for *in vivo* experiments. Significance was calculated using a 2-tailed Student's t-test (*p < 0.05, **p < 0.01, ***p < 0.001, *ns* = not significant).

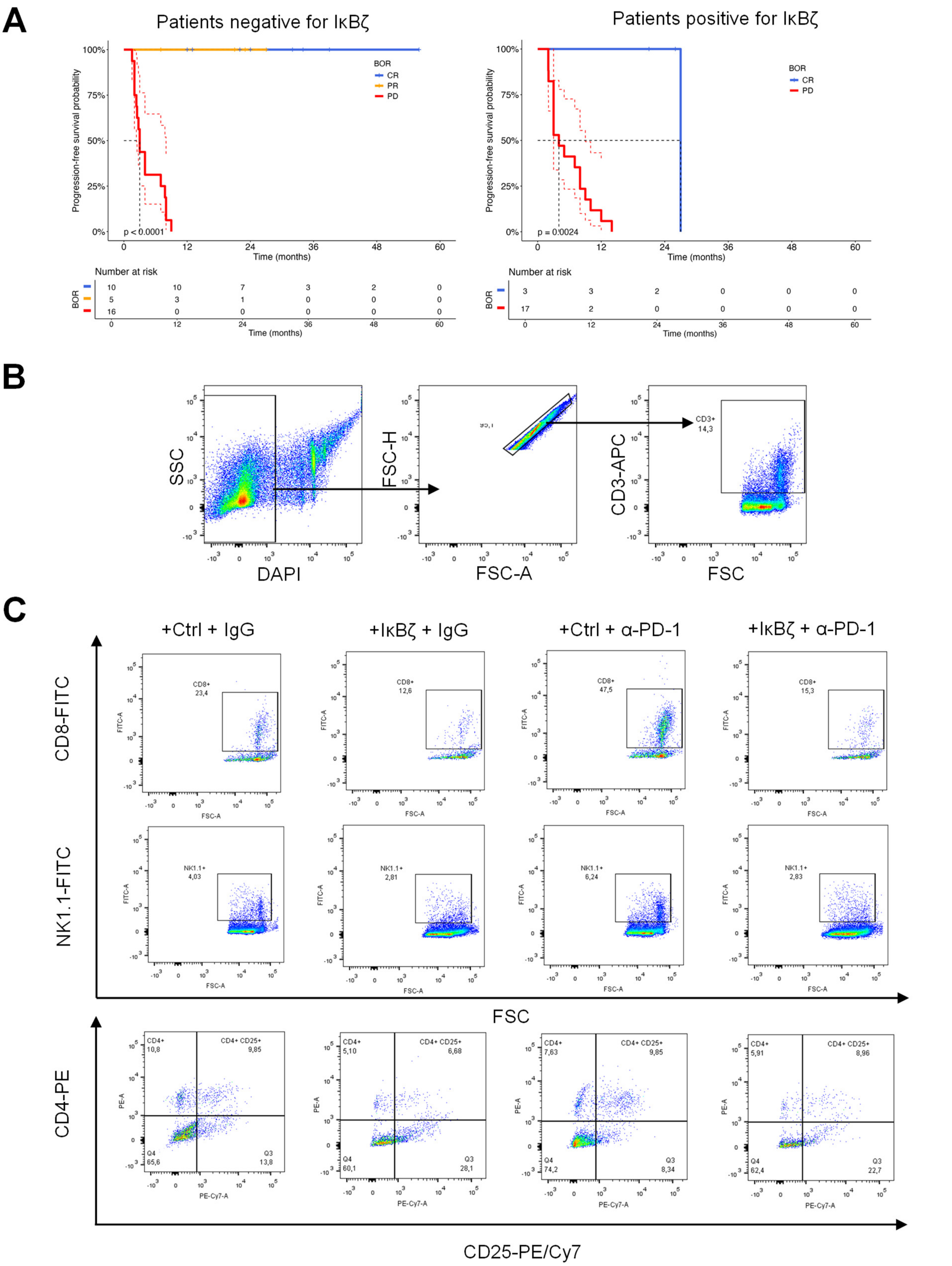


**Supplementary Figure S4. Gating strategy and representative flow cytometry plots for the analysis of α-PD-1 antibody treated control and IκBζ-overexpressing B16-F10 tumors. A.** Kaplan-Meier curve showing the progression-free survival (PFS) of immunotherapy sensitive (CR and PR) and resistant (PD) patients, grouped according to the presence or absence of tumor-derived IκBζ protein expression. Shown is the median ± 95% CI. **B.** Gating strategy used for the flow cytometry analysis. **C.** Representative plots for the flow cytometric analysis of T-cells and NK-cells in IgG- or α-PD-1-treated control and IκBζ-overexpressing B16-F10 tumors at the endpoint.

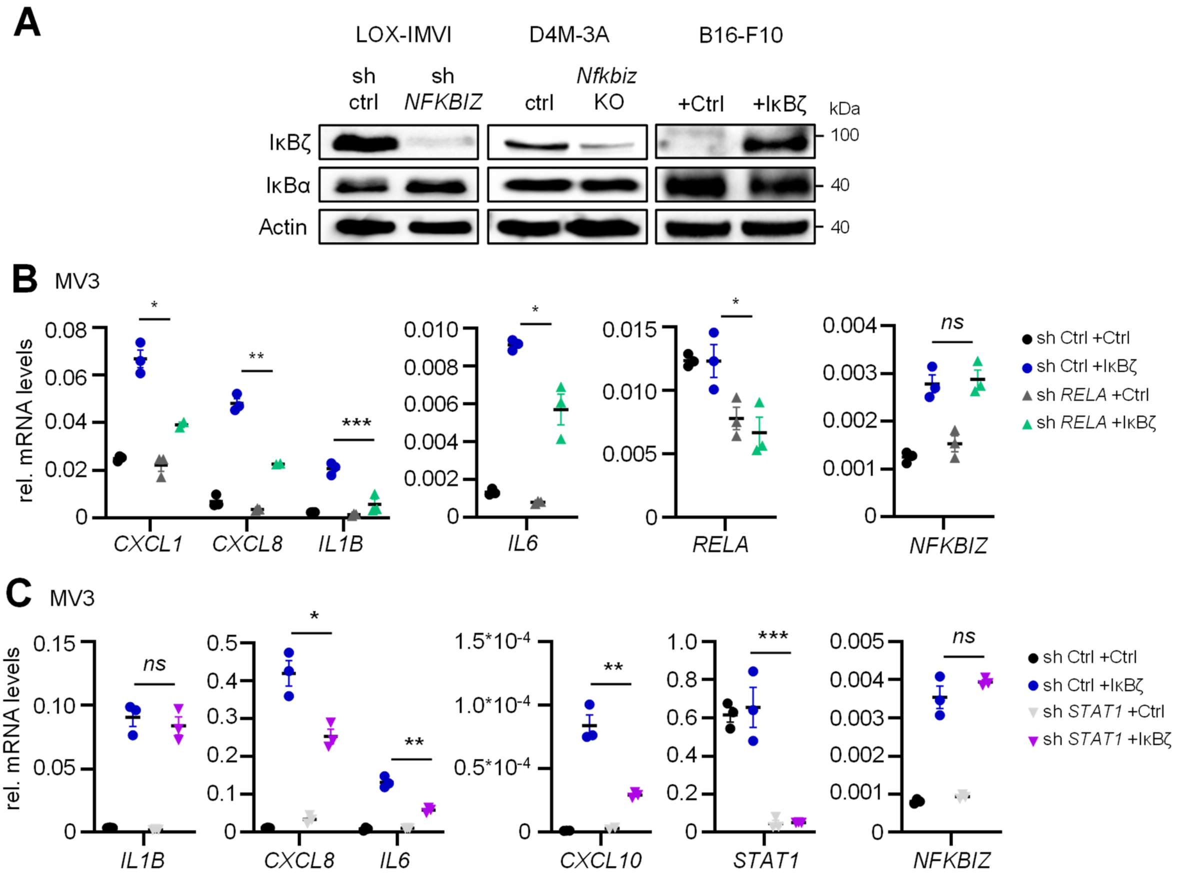


**Supplementary Figure S5. Additional data on IκBζ-dependent regulation of the transcription factor function of STAT1, STAT3, and p65. A.** Immunoblot analysis of IκBα levels in IκBζ knockdown LOX-IMVI and knockout D4M-3A cells, as well as IκBζ-overexpressing B16-F10 cells. β-actin served as a loading control. **B.+C**. Gene expression of IκBζ target genes in control or IκBζ-overexpressing MV3 cells, in the presence or absence of *RELA* (p65) (**B**) or *STAT1* (**C**). Knockdown of both genes was achieved by a lentiviral shRNA; control cells were generated using a non-coding shRNA (sh Ctrl). Subsequently, an empty plasmid or IκBζ was transiently overexpressed. Relative mRNA levels were normalized to the reference gene *RPL37A.* Shown is the mean of 3 biological replicates ± standard deviation (SD). Significance was calculated using a 2-tailed Student's t-test (*p < 0.05, **p < 0.01, ***p < 0.001, *ns* = not significant).

**Supplementary Table S1. Gene expression primer sequences.**

| Primer | Purpose (species/Ref) | Forward | Reverse |
|---|---|---|---|
| *CCL5* | human | GATCAAGACAGCACGTGGAC | TCGGGTGACAAAGACGACTG |
| *CSF3* | human | GAGGAAGATCCAGGGCGATG | AGCTTGTAGGTGGCACACTC |
| *CXCL1* | human | TCAATCCTGCATCCCCCATAG | CAGGAACAGCCACCAGTGAG |
| *CXCL8* | human | AAACTGGGTGCAGAGGGTTG | GCTTGAAGTTTCACTGGCATC |
| *CXCL9* | human | ATTGGTGCCCAGTTAGCCTC | TTCTGGCCACAGACAACCTC |
| *CXCL10* | human | TGCAAGCCAATTTTGTCCACG | CTGCATCGATTTTGCTCCCC |
| *IL1A* | human | CAGCCAGAGAGGGAGTCATTTC | CTGGAACTTTGGCCATCTTGAC |
| *IL1B* | human | TCAGCCAATCTTCATTGCTCAAG | GGTCGGAGATTCGTAGCTGG |
| *IL6* | human | CATCCTCGACGGCATCTCAG | TGCCTCTTTGCTGCTTTCAC |
| *NFKBIZ* | human | ACACCCACAAACCAACTCTGG | TGCTGAACACTGGAGGAAGTC |
| *RELA* | human | AGGCTATCAGTCAGCGCATC | AGCATTCAGGTCGTAGTCCC |
| *RPL37A* | human/Ref. gene | AGATGAAGAGACGAGCTGTGG | CTTTACCGTGACAGCGGAAG |
| *STAT1* | human | AGGTTAACGTTCGCACTCTG | GCTGCTGAAGTTCGTACCAC |
| *STAT3* | human | GACTCTCAATCCAAGGGGC | CCTCTGCCGGAGAAAGAG |
| *Actin* | mouse/Ref.gene | AGGAGTACGATGAGTCCGGC | GGTGTAAAACGCAGCTCAGTA |
| *Ccl5* | mouse | TGCTGCTTTGCCTACCTCTC | TCTTCTCTGGGTTGGCACAC |
| *Csf3* | mouse | ATCCATGGCTCAACTTTCTGC | GCTGCAGGGCCATTAGCTTC |
| *Cxcl1* | mouse | ACGTGTTGACGCTTCCCTTG | TCCTTTGAACGTCTCTGTCCC |
| *Cxcl9* | mouse | GAGCAGTGTGGAGTTCGAGG | GGCAGGTTTGATCTCCGTTC |
| *Cxcl10* | mouse | CCCACGTGTTGAGATCATTGC | CTCTGCTGTCCATCCATCGC |
| *Hprt1* | mouse/Ref. gene | CGTCGTGATTAGCGATGATGAAC | CATCTCGAGCAAGTCTTTCAGTC |
| *Il1a* | mouse | CTCATTGGCGCTTGAGTCGG | AGAGAGAGATGGTCAATGGCAG |
| *Il1b* | mouse | AGCTGAAAGCTCTCCACCTC | GCTTGGGATCCACACTCTCC |
| *Il6* | mouse | GTCCGGAGAGGAGACTTCAC | GCAAGTCGATCATCGTTGTTC |
| *Nfkbiz* | mouse | AACTCGCCAAGAGACCAGTG | AGAGCCACTGACTTGGAACG |

**Supplementary Table S2. ChIP primer sequences.**

| Primer | Purpose (ChIP) | Forward | Reverse |
|---|---|---|---|
| *CXCL1-1* | IκBζ | AGCTTCCTCCTCCCTTCTGG | AGGGCAGGAGAAGAGTGTTG |
| *CXCL1-2* | STAT3 | GATGCCCACTGCTTCTTGAC | TTAATCACGCTGCCCAAATC |
| *CXCL1-3* | P65 | AAGGCGAATATCCCAGAGTC | GCCTCGCCCTTCAGAGTAAC |
| *CXCL8* | IκBζ, STAT3, p65 | AGACAGCAGAGCACACAAGC | TCCTTCCGGTGGTTTCTTC |
| *CXCL10-1* | STAT3 | TTTGCCCTGCTCTCCCATAC | TGAGCAGGAGGACATCAGTG |
| *CXCL10-2* | IκBζ | CCATGTTGCAGACTCGAAGG | AAGAGGAGCAGAGGGAAATTC |
| *IL1B* | IκBζ, p65 | GCTAAACCAAACCCCAACTAGC | TCTCTGCCTCCCTCTCTCAG |
| *IL6* | IκBζ, STAT3, p65 | AGACATGCCAAAGTGCTGAG | TGCAGCTTAGGTCGTCATTG |
| *MB* | Negative Control | CTCATGATGCCCCTTCTTCT | GAAGGCGTCTGAGGACTTAAA |

**Supplementary Table S3. Summary of the clinical characteristics of the melanoma patients.**

| ID | BOR | IκBζ expression | PFS [months] | OS [months] | Therapy | Gender | Age | Stage |
|---|---|---|---|---|---|---|---|---|
| 1 | CR | 0 | 13 | 27 | Nivolumab + Ipilimumab | female | 52 | IV |
| 2 | CR | 0 | 22 | 83 | Nivolumab + Ipilimumab | male | 71 | IV |
| 3 | CR | 0 | 34 | 57 | Nivolumab + Ipilimumab | male | 59 | IV |
| 4 | CR | 0 | 56 | 83 | Nivolumab + Ipilimumab | female | 73 | IV |
| 5 | CR | 0 | 2 | 32 | Nivolumab + Ipilimumab | male | 88 | IV |
| 6 | PR | 0 | 5 | 38 | Nivolumab + Ipilimumab | male | 59 | IV |
| 7 | PR | 0 | 5 | 42 | Nivolumab + Ipilimumab | male | 59 | IV |
| 8 | PR | 0 | 21 | 22 | Nivolumab + Ipilimumab | male | 81 | IV |
| 9 | PR | 0 | 27 | 63 | Nivolumab + Ipilimumab | male | 85 | IV |
| 10 | PD | 0 | 23 | 60 | Nivolumab + Ipilimumab | male | 84 | IV |
| 11 | PD | 0 | 3 | 7 | Nivolumab + Ipilimumab | female | 81 | IV |
| 12 | CR | 0 | 32 | 32 | Nivolumab | male | 75 | IV |
| 13 | CR | 0 | 34 | 34 | Nivolumab | female | 85 | IV |
| 14 | CR | 0 | 56 | 56 | Nivolumab | female | 74 | IV |
| 15 | CR | 0 | 39 | 39 | Nivolumab | male | 89 | IV |
| 16 | PD | 0 | 8 | 14 | Nivolumab + Ipilimumab | male | 74 | IV |
| 17 | PD | 0 | 9 | 13 | Nivolumab + Ipilimumab | male | 55 | IV |
| 18 | PD | 0 | 2 | 18 | Nivolumab + Ipilimumab | male | 68 | IV |
| 19 | PD | 0 | 7 | 21 | Nivolumab + Ipilimumab | female | 56 | IV |
| 20 | PD | 0 | 8 | 23 | Nivolumab + Ipilimumab | male | 61 | IV |
| 21 | PD | 0 | 2 | 10 | Nivolumab | male | 79 | IV |
| 22 | PD | 0 | 4 | 6 | Nivolumab | male | 94 | IV |
| 23 | PD | 0 | 2 | 4 | Nivolumab | female | 85 | IV |
| 24 | PD | 0 | 3 | 10 | Nivolumab + Ipilimumab | female | 65 | IV |
| 25 | PD | 0 | 4 | 6 | Nivolumab + Ipilimumab | female | 57 | IV |
| 26 | CR | 0 | 24 | 30 | Nivolumab + Ipilimumab | male | 84 | IV |
| 27 | PD | 0 | 2 | 4 | Nivolumab + Ipilimumab | female | 66 | IV |
| 28 | PD | 0 | 3 | 13 | Nivolumab + Ipilimumab | female | 59 | IV |
| 29 | PD | 0 | 8 | 27 | Pembrolizumab | female | 39 | IV |
| 30 | PD | 0 | 2 | 4 | Nivolumab | male | 76 | IV |
| 31 | PD | 0 | 2 | 8 | Pembrolizumab | male | 68 | IV |
| 32 | PD | 1 | 3 | 11 | Nivolumab + Ipilimumab | male | 67 | IV |
| 33 | CR | 1 | 21 | 27 | Nivolumab + Ipilimumab | male | 64 | IV |
| 34 | CR | 1 | 27 | 27 | Nivolumab | female | 86 | IV |
| 35 | PD | 1 | 8 | 24 | Nivolumab + Ipilimumab | female | 36 | IV |
| 36 | PD | 1 | 12 | 15 | Nivolumab + Ipilimumab | male | 77 | IV |
| 37 | PD | 1 | 3 | 11 | Nivolumab + Ipilimumab | male | 69 | IV |
| 38 | PD | 1 | 2 | 105 | Pembrolizumab | male | 48 | IIIC |
| 39 | PD | 1.5 | 3 | 7 | Nivolumab + Ipilimumab | male | 59 | IV |
| 40 | PD | 1.5 | 14 | 15 | Nivolumab | female | 88 | IV |
| 41 | PD | 1.5 | 8 | 14 | Nivolumab | male | 76 | IV |
| 42 | PD | 2 | 2 | 9 | Nivolumab + Ipilimumab | female | 62 | IV |
| 43 | PD | 2 | 5 | 10 | Nivolumab + Ipilimumab | female | 56 | IV |
| 44 | PD | 2 | 10 | 18 | Nivolumab + Ipilimumab | male | 69 | IV |
| 45 | PD | 2 | 3 | 8 | Nivolumab | male | 82 | IV |
| 46 | PD | 2 | 4 | 8 | Nivolumab + Ipilimumab | male | 67 | IV |
| 47 | PD | 2 | 3 | 17 | Nivolumab | male | 81 | IV |
| 48 | CR | 2 | 26 | 77 | Nivolumab + Ipilimumab | male | 56 | IV |
| 49 | PD | 2 | 7 | 66 | Nivolumab + Ipilimumab | male | 51 | IV |
| 50 | PD | 2 | 2 | 31 | Nivolumab + Ipilimumab | male | 57 | IV |
| 51 | PD | 2 | 9 | 19 | Nivolumab | male | 88 | IV |
